# Osmolyte homeostasis controls single-cell growth rate and maximum cell size of *Saccharomyces cerevisiae*

**DOI:** 10.1101/586479

**Authors:** Tom Altenburg, Björn Goldenbogen, Jannis Uhlendorf, Edda Klipp

## Abstract

Cell growth is well described at the population level, but precisely how nutrient and water uptake and cell wall expansion drive the growth of single cells is poorly understood. Supported by measurements of single-cell growth trajectories and cell wall elasticity, we present a single-cell growth model for yeast. The model links the thermodynamic quantities turgor pressure, osmolarity, cell wall elasto-plasticity, and cell size, using concepts from rheology and thin shell theory. It reproduces cell size dynamics during single-cell growth, budding, and hyper- or hypoosmotic stress. We find that single-cell growth rate and final size are primarily governed by osmolyte uptake and consumption, while bud expansion depends additionally on different cell wall extensibilities of mother and bud. Based on first principles the model provides a more accurate description of size dynamics than previous attempts and its analytical simplification allows for easy combination with models for other cell processes.

## Introduction

Cells are exposed to hydrostatic pressure driven by the difference between inner and outer osmolarity. At the boundaries of tissues in higher eukaryotes, cells often encounter a wide range of osmotic changes due to cytokines, hormones etc. Similarly, unicellular organisms such as the eukaryotic model organism *Saccharomyces cerevisiae* or yeast, proliferate under a wide range of osmotic conditions, caused for example by periods of rain or drought. In the presence of these changing conditions yeast has evolved strategies to maintain cellular integrity, ranging from regulating intracellular osmolarity to constructing elastic scaffolds such as the cytoskeleton or the cell wall. Water flow over the cell membrane tends to equilibrate osmotic and hydrostatic pressure differences (1) and, therefore, impacts cell size, depending on cellular deformability. Therefore, while on the one hand yeast has to adapt its internal osmotic pressure to external conditions (2–4) to prevent bursting, as well as critical shrinking, it also has to regulate growth rate, depending on nutrient uptake and subsequent metabolization. Nutrient uptake not only provides building blocks and energy for the synthesis of new cell material, but also increases internal osmolarity and thereby drives inward water flux, which in turn leads to an increase in cell size. In walled cells, such as bakers yeast or plant cells, the difference between internal and external osmotic pressure is counteracted by turgor pressure arising from elastic expansion of cell wall material. Turgor pressure prevents exaggerated swelling and maintains cell shape. Although reported values of turgor pressure in yeast range from 0.1 MPa to 1 MPa (5, 6), more recent single-cell measurements suggested a value of 0.2 MPa (7).

While various studies have attempted to address aspects of the impacts of osmo-regulation on single cell growth these issues remain poorly understood. In a previous model, thermodynamic descriptions of volume and pressure changes were integrated within the osmotic stress response system, i.e. the high osmolarity glycerol (HOG) signaling pathway, metabolism and gene expression (3). This integrative model permitted predictions regarding the effect of several gene-knockouts on volume dynamics. Another model integrated further published data with biophysical and mechanical properties of yeast to describe the loss in volume immediately after osmotic stress (4). Both models explain volume regulation following a hyperosmotic shock, but are not designed to describe the small and steady volume variations during normal growth.

Even though various volume regulation models have been proposed, a unified understanding of the interplay between cell mechanics, turgor, volume, and metabolism during growth and perturbations, e.g. osmotic shocks, is still missing. Existing approaches addressing these issues focus solely on animal cells, where cellular integrity is maintained by the cytoskeleton (8, 9). However, mammalian cells also face high osmotic pressure changes and cell integrity of certain species is supported by external structures such as matrix, mucus or wax, which fulfill the same function as a cell wall.

Here, we present a single-cell growth model (SCGM), which focuses on the interplay of three thermodynamic quantities: cell volume, osmolarity and turgor pressure, and which covers growth and budding of single yeast cells as well as the response to external osmotic variations. We further tested the model against single-cell growth data from brightfield microscopy images and used atomic force microscopy (AFM) to gain information on the cell wall elasticity during budding. The model combines different concepts, such as cell wall mechanics in yeast (10–13) (14, 15), rheology, a subfield of continuum mechanics and broadly used in plant physiology (16–19) and applied to fungi (20, 21), thin shell theory (22–24), water homeostasis and dynamics (1, 25) and osmoregulation (in general or exemplified by HOG) (3, 26, 27). The SCGM is capable of describing both drastic volume variations caused by hyper- or hypoosmotic shocks, as well as relatively small but steady gains in cell size due to growth. To demonstrate that the SCGM can be combined with models for cellular signaling and metabolism, we introduced the HOG signaling cascade model (27) as an exemplary pathway that plays a major role in yeast osmoregulation (2).

## Results

### The single-cell growth model combines formalisms for turgor pressure, osmo-regulation and cell wall mechanics

Cellular volume varies according to material accumulation and water flux across the cell membrane, which follows the osmotic and hydrostatic pressure gradient. For volume flux and the conversion from osmolarity to osmotic pressure we considered established formalisms described by Kedem-Katchalsky and Boyle van’t Hoff (1, 3, 25, 27). To this end, we defined total cell size *V*_*t*_ = *V*_*os*_ + *V*_*b*_ as the sum of osmotic volume *V*_*os*_, which is sensitive to changes in osmotic or hydrostatic pressures, and solid volume *V*_*b*_, which is not affected by water dynamics (predominantly macro-molecules and organelles contribute to *V*_*b*_), see Fig. 1A. In a confined system, such as a cell, the outward-directed water flux over the boundary, the cytoplasmic membrane, must equal the negative change in volume of this system over time:

**Fig. 1.**
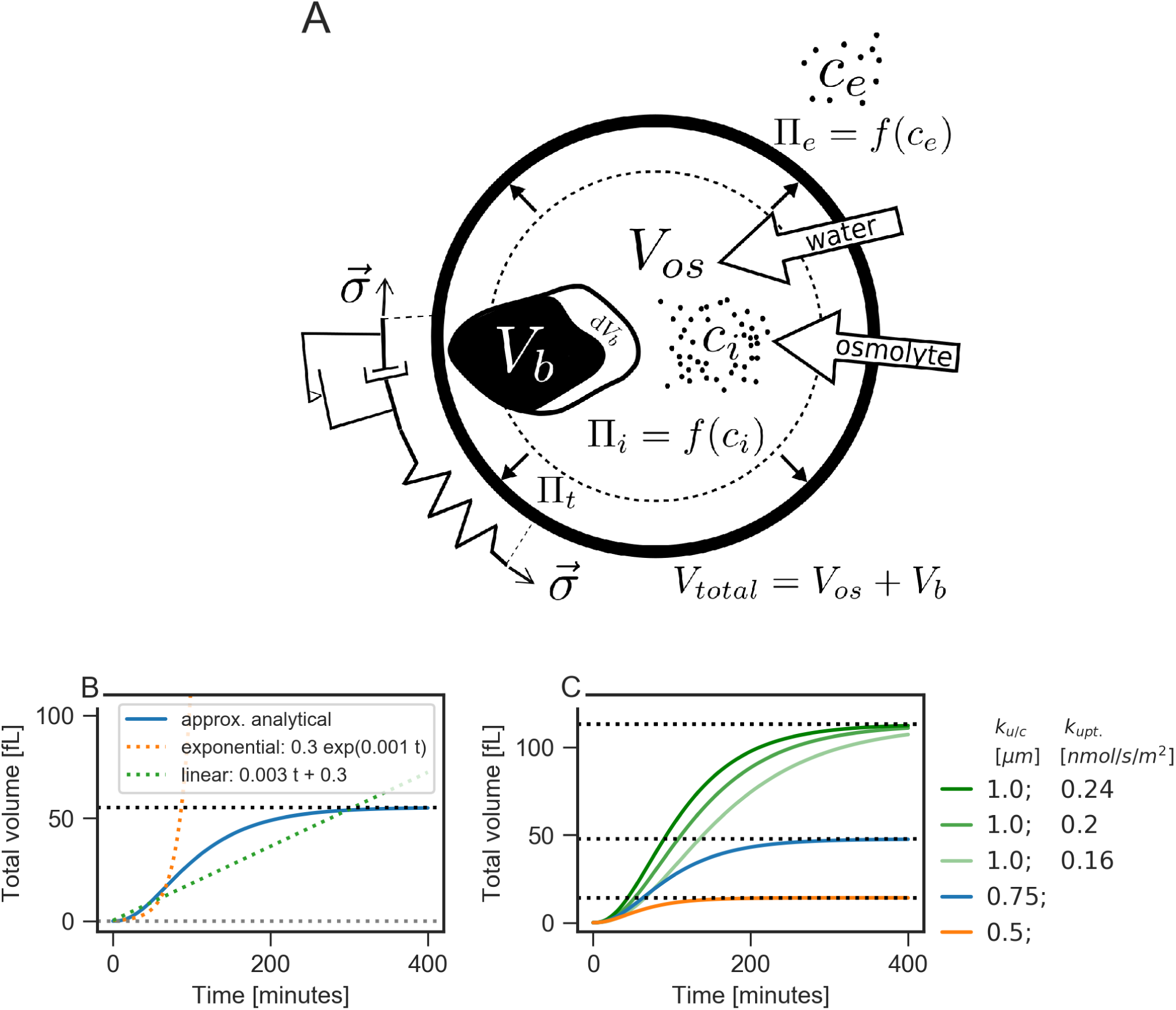
The single-cell growth model (SCGM) predicts non-linear growth behavoir. **A** SCGM sketch. **B and C** Volume trajectory for an individual yeast cell. **B** The SCGM (blue line) shows different volume trajectories compared to linear (green dotted line) or exponential (orange dotted line) growth. **C** Varying uptake and consumption equally (*k*_u/c_ = *const*.), leaves final cell size *V*_final_(*t* → ∞) unaffected, but changes growth rate (green lines). Alterations of ratio *k*_u/c_ affects final cell size *V*_final_ with *V*_final_ = 12*π*(*k*_u/c_)^3^. SCGM is, as antimony model, available under: https://github.com/tbphu/volume_model/blob/master/volume_reference_radius.txt

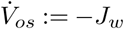

According to Kedem-Katchalsky, water flux *J*_*w*_ a cross the membrane is proportional to the causative force, i.e. pressure differences, and is thus defined as:

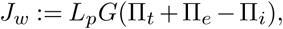

where *L*_*p*_ denotes the hydraulic conductivity of the membrane per unit area, *G* is the area of the cell surface, Π_*t*_ is the turgor pressure and Π_*e*_ and Π_*i*_ are the external and internal osmotic pressures. Turgor pressure is normally calculated under a quasi-steady state assumption of negligible water fluxes (*J*_*w*_ = 0) and, hence, equals the difference between external and internal osmotic pressure. To capture the dynamics of volume variation, we went a step further by neither constraining *J*_*w*_ nor the total number of particles in the system, thereby establishing distinct mathematical descriptions for the three main quantities: volume, turgor pressure, and osmolarity. Uptake and dilution of the osmotically active compounds, such as nutrients and ions, determine the internal and external osmotic pressures according to Boyle van’t Hoff’s equation:

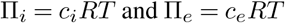

where *c*_*i*_ and *c*_*e*_ are the internal and external osmolyte concentrations, *R* is the gas constant and *T* the temperature. Osmolyte uptake *ċ*_*i*_ was assumed to be proportional to the cell surface, over which osmolytes must be transported, while consumption of osmolytes was assumed to be proportional to the cell volume. Additionally, volume expansion must lead to a dilution of osmolytes, hence to a decrease in *c*_*i*_. *k*_uptake_ and *k*_consumption_ are the respective proportionality or rate constants. Combined, we obtain the following simplified description of osmolyte dynamics within the cell:

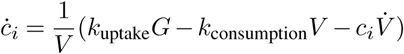

This basic description can be extended if a more detailed metabolism model is at hand, as we exemplify below for osmotic stress response by including the HOG pathway and glycerol accumulation.

Having derived descriptions for volume and inner osmolarity, we can describe turgor pressure in the following (for details see Supplementary information). Given that turgor pressure equals hydrostatic pressure acting on the cell wall, the main structural element of the yeast cell, the cell wall and its mechanical properties must be considered in our description. Note, for simplification, we assumed the cell wall to be a thin spherical shell with radius *r*. The cell wall expands elastically or plastically depending inner pressure forces. While an elastically expanded volume relaxes to its initial size when pressure vanishes, plastic expansion is irreversible. Taking a constant wall thickness into account the irreversible expansion can also been interpreted as cell growth, where new cell wall material has to be provided by the cell wall synthesis machinery. The Hookean element represents the elastic response, while the Bingham element represents the plastic response. Assuming a linear constitutive relationship for elasticity, the deformation of the cell volume, i.e. strain *ε*_*Hook*_, scales linearly with turgor pressure in a purely elastic cell wall

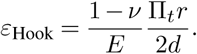

Here *E* is Young’s modulus representing the elasticity of the cell wall, *v* is Poisson’s ratio, and *d* is thickness of the cell wall. Consequently, water influx will lead to higher stress in the cell wall and directly to larger strains. Note that cell wall thickness is assumed to be constant and therefore depended neither on the radius *r* nor on the time *t*.

In contrast to elastic deformation, plastic deformation, was assumed to occur only above the critical turgor pressure Π_*tc*_ and instead of the strain, the strain rate is proportional to the acting pressure with a factor *ϕ*. Whereby *ϕ* represents the extensibility of the cell wall. The Bingham strain rate reads:

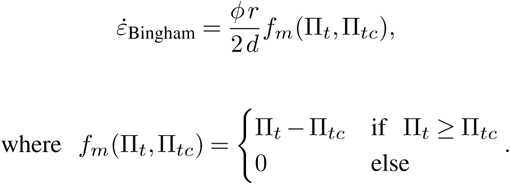

To reflect elastic and plastic behavior of the cell wall, we used a mechanical model of the cell wall in which a Hookean and a Bingham element are coupled in series Fig. 1 A. Here, total strain is the sum of individual strains of each element 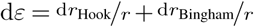. Taking the time derivative 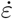 of the total strain enabled us to combine descriptions for both mechanical elements. Solving the resulting equation (see Supplementary Material) for the change in turgor pressure over time 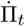 we derived the following ODE for turgor pressure:

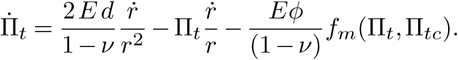

Models that share two of the three terms of this equation have already been applied in other turgor-related models of (16, 17, 19, 28). In a comparison with the previously proposed description of turgor in plant physiology (17) showed that the additional term resulted from geometric description of the cell as a thin spherical shell, in particular when calculating the time derivative of the elastic element (see Supplementary Material).

Cell expansion is therefore mainly influenced by two processes: first, the control of internal osmolarity *c*_*i*_ by uptake, dilution and consumption of osmolytes, which together with turgor pressure Π_*t*_ drive water influx and outflux *J*_*w*_ and hence determine water volume in the cell (*V*_*os*_), and second, elasto-plastic deformation of the cell wall due to turgor pressure. The water influx *J*_*w*_ *<* 0 expands the cell wall elastically and thereby increases turgor pressure. When turgor pressure exceeds the critical value Π_*tc*_, the cell wall starts to yield, e.g. the cell wall expands irreversibly and the cell grows. Yielding leads to relaxation of stress in the cell wall and in turn to a decrease in turgor pressure, facilitating further influx of water. Hence, growth typically proceeds close to, but not at steady state (*J*_*w*_ = 0) where water and osmolyte influxes are balanced by turgor dynamics.

Summarizing, the basic SCGM comprises three coupled ODE’s for volume, turgor pressure and internal osmolarity. Key parameters and initial values for simulation or analytic solution are listed in Tab. 1. The temperature was set to T = 303 *K*, i.e. the optimal growth temperature for yeast and our experimental standard condition. The external osmolyte concentration *c*_*e*_ was set to 240 mM, the mean value for our standard growth medium.

**Table 1.** Parameters and initial values.

| parameter | value | unit | description | source |
| --- | --- | --- | --- | --- |
| $r_b^0$ | 0.3 | $\mu\text{m}$ | init. radius (non-osmolytic volume) | assumption |
| $r_{os}^0$ | 0.1 | $\mu\text{m}$ | init. radius (osmolytic volume) | assumption |
| $c_i^0$ | 319.17 | mM | osmolyte concentration (internal) | calculated from turgor pressure |
| $c_e$ | 240.0 | mM | osmolyte concentration (external) | measured |
| $\Pi_t^0$ | $2.0 \times 10^5$ | Pa | initial turgor pressure | (7) |
| $R$ | 8.314 | $\frac{\text{J}}{\text{mol K}}$ | ideal gas constant | |
| $T$ | 303.0 | K | temperature | |
| $Lp$ | $1.19 \times 10^{-6}$ | $\frac{\mu\text{m}}{\text{s Pa}}$ | membrane water permeability | (3) |
| $\Pi_{tc}$ | $2.0 \times 10^5$ | Pa | critical turgor pressure | (7) |
| $d$ | 0.115 | $\mu\text{m}$ | cell wall thickness | (29) |
| $\phi$ | $1.0 \times 10^{-3}$ | $\frac{1}{\text{s Pa}}$ | extensibility | initial assumption, fit in Fig. 4 |
| $\nu$ | 0.5 | — | Poisson's ratio | |
| $E$ | $2.58 \times 10^6$ | Pa | Young's modulus (3D) | (7) |
| $k_{\text{uptake}}$ | $2.0 \times 10^{-16}$ | $\frac{\text{mmol}}{\mu\text{m}^2 \text{s}}$ | osmolyte uptake rate constant | initial assumption, fit in Fig. 4 |
| $k_{\text{consumption}}$ | $2.0 \times 10^{-16}$ | $\frac{\text{mmol}}{\mu\text{m}^3 \text{s}}$ | osmolyte consumption rate const. | initial assumption, fit in Fig. 4 |

Considering a cell growing without any osmotic perturbations, the SCGM can be reduced to a simpler description of cell size development. Simulations revealed that without osmotic stress, internal osmolarity *c*_*i*_ and turgor pressure Π_*t*_ approach steady states. In this case the SCGM can be solved analytically (Fig. 1A, Fig. S7). We obtained an implicit differential equation *F* comprising the time derivative of the radius 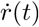, the time-dependent function of cellular radius *r*(*t*), steady states and time:

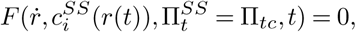

where 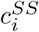 is a quasi-steady state for internal osmolarity (quasi-, because still depends on r(t)) and 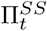 the steady state for turgor pressure, which equals critical turgor pressure Π_*tc*_ during growth. Solving F (see Supplementary Material), we first, identified a limes of cellular radius for long times scales:

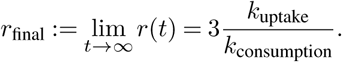

and second, obtained an analytical solution for *r*(*t*). The resulting expression for *r*(*t*) is rather complex (see Supplementary Material). Therefore, we approximated 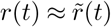, yielding a function for the radius of a growing cell:

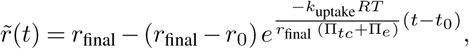

where *r*_*final*_ is the final cellular radius for long time scales *t, r*_0_ is the initial radius at time *t*_0_. The solution points out that external osmotic pressure, critical turgor pressure and internal osmotic pressure, as well as *k*_uptake_ and *k*_consumption_, dictate the trajectory of cell growth. For our solution it was crucial to include the quasi-steady state 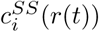. In this way, the influences of *k*_uptake_ and *k*_consumption_ on the quasisteady state of internal osmolarity could be propagated to the radius description 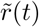. As shown in Fig. 1C, our model suggests that final cell size depends exclusively on the ratio *k*_u/c_ = *k*_uptake_*/k*_consumption_, but not on cell wall-related quantities. Scaling both rate constants equally while keeping ratio *k*_u/c_ constant, scales growth rate, but keeps final cell size constant. The growth dynamics 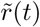 depends additionally on critical turgor pressure and external osmolarity. For hyperosmotic shock response, we derived another analytical solution for turgor dynamics, which is related to the Merritt-Weinhaus equation (30), i.e. the pressure-to-size relation of an ideal elastic thin shell (Fig. S8).

### Coupled SCGM permits the description of combined growth of mother and bud

Above, we have provided a description of growth dynamics for an individual cell, which is only valid for yeast cells in G1 phase (single cell without a bud). In other cell cycle phases, cells are combined of two spherical compartments (mother and bud) with different growth rates (31), and are connected by a neck allowing free exchange of osmolytes and water. Therefore, we coupled two SCGMs, one each for mother and bud (Fig. 2A), in which both start with the same small initial volume, but with a time delay *t*_budstart_ for the bud, leaving time for the mother to grow. Hence, *t*_budstart_ represents characteristic cell cycle length, e.g. the time between two successive bud formations. The instances were coupled by allowing water flux according to pressure differences and diffusion of *c*_*i*_ between the compartments. The coefficients for both fluxes were arbitrarily chosen such that water and osmolyte gradients vanish in the considered time scale. Except for mechanical cell wall properties and initial geometry, both instances were similar with equal *k*_uptake_ and *k*_consumption_. The assumption of similar mechanical properties of the cell wall for both compartments is very unlikely, since alterations of the cell wall structure during budding were reported (32). In particular, chitin incorporation into the lateral cell wall is delayed in buds until after septation. As potentially discriminating cell wall property, we considered either the Young’s modulus *E* or the extensibility *ϕ*. Systematic analysis of the impact of varying *E* or *ϕ* on bud growth revealed that only a strongly decreased *E*_bud_ compared to *E*_mother_ led to bud growth (Fig. S9), though at the expense of mother growth. In contrast, when varying *ϕ*, both compartments continued to grow (Fig. 2B), as long as *ϕ*_bud_ was at least 10^1.5^ times higher than *ϕ*_mother_. Note that we focused here on biophysical principles, instead of on the complex biochemical processes governing cell division, such as cell polarization and subsequent bud emergence.

**Fig. 2.**
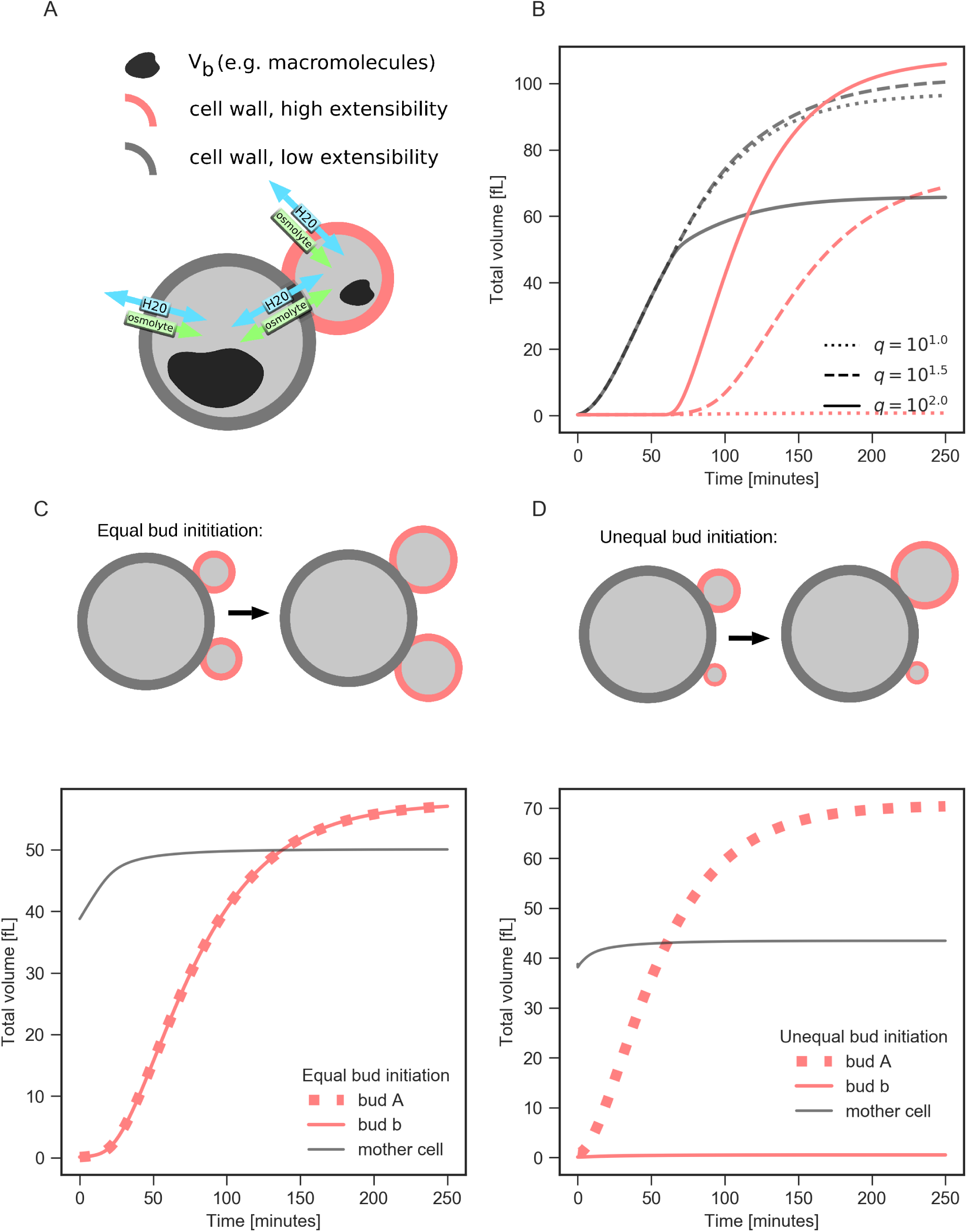
Bud expansion driven by cell wall anisotropy and a possible protection mechanism against two-bud growing malfunction. **A** Two volume model instances were interconnected via water and osmolytes fluxes. The coupled model accounts for different cell wall extensibilities *ϕ* between bud and mother. **B** Volume dynamics are shown for mother and bud for three different extensibility ratios *q* = *ϕ*_bud_ */ϕ*_mother_. Bud expansion is possible only at *q >* 10. **C** To investigate two-bud growing mutants, two bud instances linked to a mother cell were initialized with equal bud sizes. Both buds have identical volume dynamics. **D** Effect of unequal initial bud sizes: differences are amplified and only the larger bud grows. The coupled SCGM is available as antimony model under: https://github.com/tbphu/volume_model/blob/SCGM/volume_mother_and_bud.txt

### Measurements of local cell wall elasticity of mother and bud reveal a distinct difference between two asymmetric growth processes in yeast, budding and shmooing

As discussed above, a significantly lower elastic modulus at the bud cell wall could drive bud expansion in the model. Earlier, we reported that such localized softening of cell wall material occurs during sexual conjugation in *S. cerevisiae* (7). To test whether both processes, budding and sexual conjugation, follow similar underlying principles, we applied multi-parametric atomic force microscopy (AFM) on living yeast cells as depicted in Fig. 3A. From the force-response curves of nano-indentation measurements (Fig. 3B), we obtained the local Young’s modulus *E*. To compare mother and bud, we selected regions of equal size at each compartment (Fig. 3C,D). We avoided strongly tilted regions, compared to the scanning plane, and regions of previous budding events, so-called bud scars, which contain a higher amount of chitin and are reported to be stiffer (33). Intriguingly, bud cell walls appeared to be stiffer, not softer, than their mothers’ cell walls (Fig. 3). From a linear fit, we estimated a 1.3 ± 0.1-fold increase in the Young’s modulus from mother to bud (Fig. S11). Although the apparent stiffness at the bud cell wall could be biased by different surface curvatures, a significant cell wall softening at the bud, as measured for sexual conjugation, can be rejected. This reveals a distinct difference between two of the asymmetric growth processes in yeast, budding and sexual conjugation. Consequently, we focused on the extensibility as distinguishing feature between mother and bud cell wall.

**Fig. 3.**
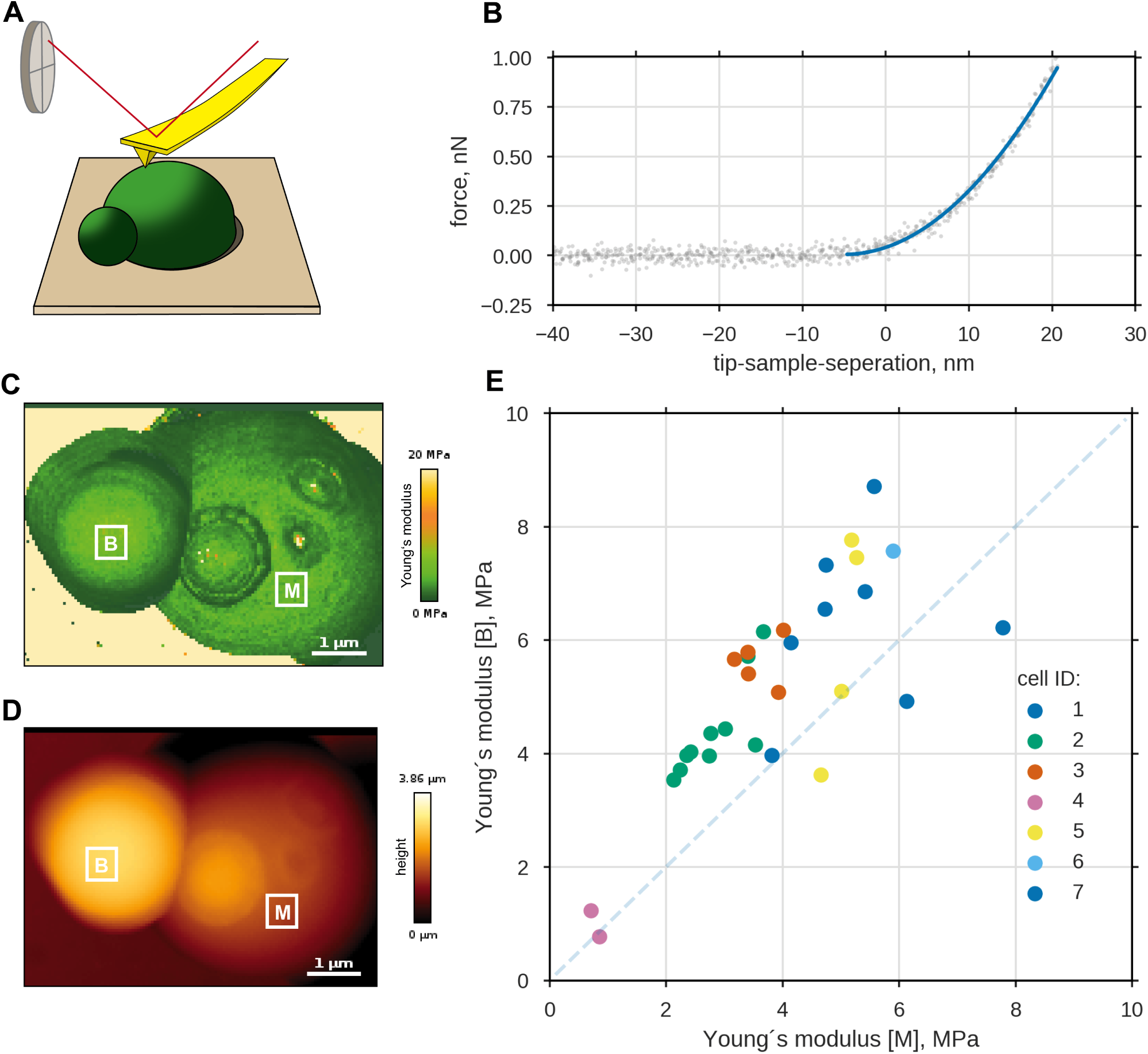
Local Young’s modulus of the cell wall is slightly higher for bud than for mother. **A** Experimental setup: entrapped haploid *S. cerevisiae* cells were scanned using atomic force microscopy, whereby at each image position the force response to nano-indentation was measured. **B** Exemplary approach curve for one pixel along with a Sneddon fit (blue). Obtained spatial information is shown for cell wall elasticity **C** and height **D** of a budding mother cell. For further analyses, the mean cell wall elasticities from the least curved region of mother and bud ([M] and [B]) were used, avoiding regions of former budding events, bud-scars. **E** Mean local *E* at buds compared to mean local *E* at mothers for 30 measurements on seven cells (same cells, same color). Dashed line: equal cell wall elasticities of bud and mother.

### Verification of the model against volume data from light microscopy

To test whether the coupled SCGM correctly describes the growth of single yeast cells and to estimate otherwise inaccessible parameters, we measured the growth trajectories of single yeast cells and used these to constrain model parameters. Cells were grown as monolayer in a microfluidic device, allowing observation of individual cells for long time periods (> 15 h). We tracked single cells in microscopic bright field images using the software CellStar (34) and manually determined the first bud emergence for each cell (Fig. 4A). A bud neck marker (Cdc10-mKate2) was used to identify pairs of mothers and corresponding buds and to reconstruct the lineage (see Methods section). Knowing the lineage enabled us to collect 21 coupled volume traces for mother and bud. Since the time between first appearance of a cell and its first budding event varied greatly, we did not fit the model to mean growth data, but to different single cell traces instead. Fig. 4B and Fig. S12 show that the coupled SCGM is able to describe the growth pattern of single cells growing at different rates. For both traces, mother and bud, six independent parameters (mean ± sd) have been determined, presented in Fig. 4C. In addition to the individual parameters initial radius *r*_0_ and time of bud appearance, *t*_budstart_, we considered two global parameters, *k*_uptake_ and *k*_u/c_ and two compartment-dependent parameters *ϕ*_mother_ and *q*=*ϕ*_bud_*/ϕ*_mother_. *r*_0_ was the fitted radius belonging to the initial osmotic volume of the later mother. With 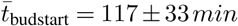 the estimated mean cell cycle length differed remarkably from the time when a new bud was clearly recognizable for the first time (first green data point). When analyzing estimated rates constants for osmolyte uptake and consumption, we found that the growth rate determining *k*_uptake_ showed some variation between single cells (1.59 × 10^*-*16^ ± 0.72 × 10^-16^ *mmol μm*^-2^ *s*^-1^). In contrast *k*_u/c_, which limits the maximum volume, showed very little variation (0.82 ± 0.07 *μm*^-1^). Focusing on cell wall extensibility, we estimated a value of 0.57 ± 0.34 *kPa*^-1^ *s*^-1^ for the mother *ϕ*_mother_ and found that *ϕ*_bud_ needs to be at least 100 times higher. We could not determine a precise value for the extensibility ratio *ϕ*_bud_*/ϕ*_mother_. However, profile likelihood and sensitivity analysis (Fig. S13) revealed that variation of *ϕ*_bud_*/ϕ*_mother_ has no significant impact on other parameters.

**Fig. 4.**
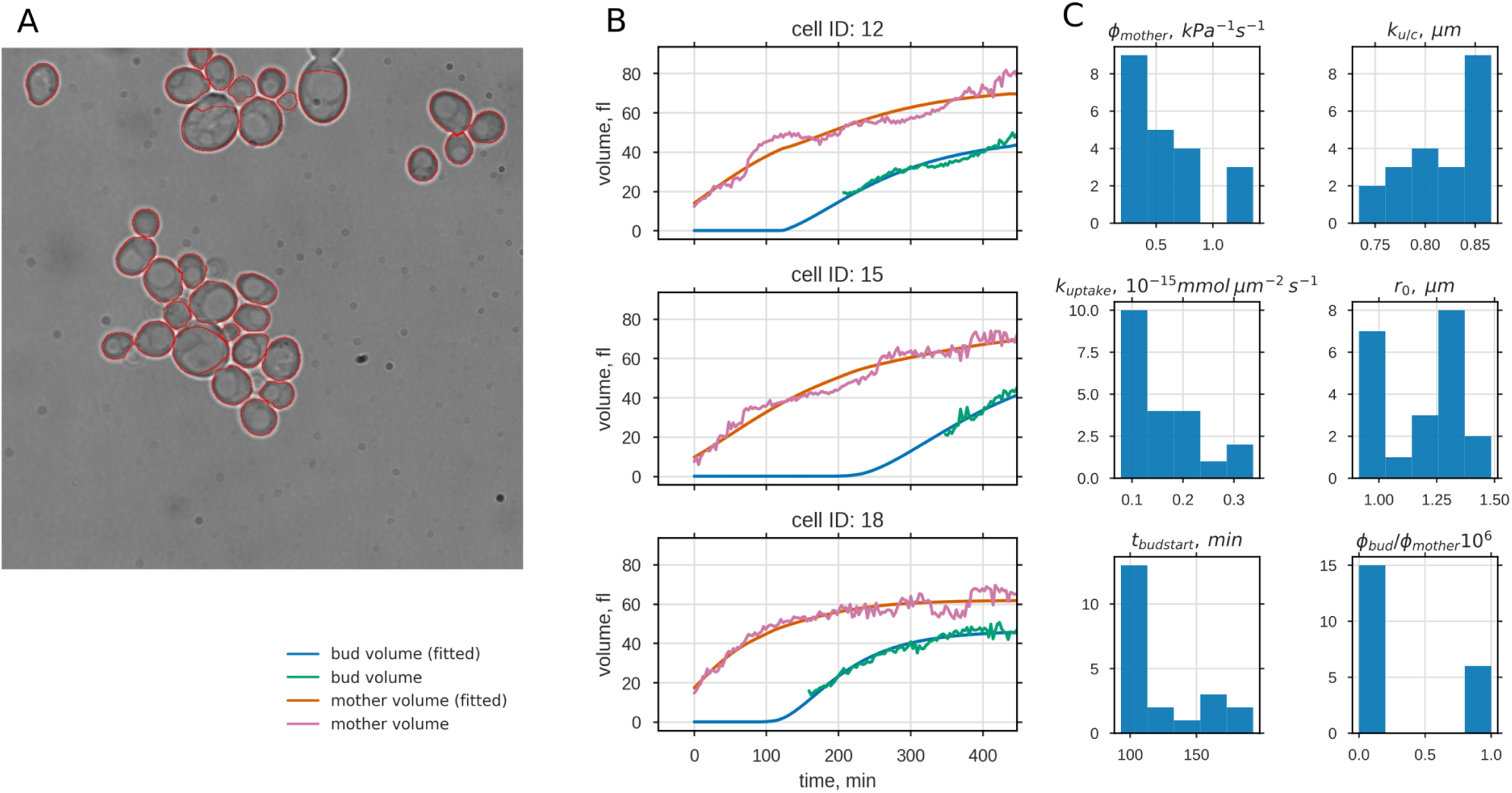
Growth parameter estimation from time series of bright field microscopy images. **A** Cross-sectional areas of 21 single yeast cells were automatically detected from transmission microscopy images and followed over time. Estimated volume increases of mother and bud were used to fit the model parameters: import rate of osmolytes *k*_uptake_, import and degradation ratio of osmolytes *k*_u/c_, cell wall extensibility of the mother *ϕ*_mother_, and the extensibility ratio of bud and mother *ϕ*_bud_ */ϕ*_mother_. Time of bud start *t*_budstart_ and initial osmotic radius *r*_0_ were used as free initial conditions. Three exemplary sets of data and corresponding fits are shown in **B** (for the complete sets see Fig. S12). Histograms of the resulting parameter values are shown in **C**. Except for *ϕ*_bud_ */ϕ*_mother_ we could assign all of the parameters with low variability, despite the high variability in the volume development in these data.

For the Δ*cdc42* mutant a probability for the emergence of two buds was reported. Our model suggests that this probability (occurrence of multiple buds) is not only influenced biochemically (e.g. by Cdc42p) but also by the the passive dynamics in cell size described above. Therefore, the constellation of two small volumes connected to a main volume establishes an additional layer of filtering out stunted buds.

### Analysis of volume trajectories from Garmendia-Torres et al. (35)

So far the analysis of growth parameters was based on 21 volume-trajectory pairs, which provided valuable information on the dimension of each parameter. However, for the extensibility ratio *ϕ*_bud_*/ϕ*_mother_ only a lower boundary could be determined and the relative small sample size did not allow for decisive statements on the parameter distribution. To increase the sample size and further challenge the cSCGM we searched for single-cell data with time resolved volume information for mother and bud and found a recent study by Garmendia-Torres et al. (35).

They reported a new microscopy-based method to investigate yeast cell-cycle and cell-size progression in parallel, by monitoring volume and histone levels of individual yeast cells. Using an automatized experimental setup they followed up to 15.000 cell cycles for each of the investigated 22 cellcycle related mutants. This innovative approach of tracking the fluorescence of fast maturating HTB2-sfGFP fusion proteins over time and assigning its intensity to distinct cell cycle phases, allowed them to relate the volumes of mother and bud at certain cell cycle stages to the time spend in each phase. Although this analytic approach revealed intriguing relations between cell cycle and cell size, it simplifies the dynamics of the cell size progression in the data. In particular, they assumed linear growth during the budded and unbudded phase. To shed light on this cell-size dynamics and to further challenge the cSCGM, the model was fitted to the experimental data (hereinafter also referred to as *data 2*) provided by Gilles Charvin. This allowed the comparison between completely independent data sets from different laboratories and the analysis of data with a drastically increased the sample size.

#### Data and data selection

The advantage of this data set, besides its dimension, is the resemblance of the used experimental and analytic approach to the approach we used. In particular, yeast strain background and culture medium, BY4741 and synthetic medium supplemented by amino acids and glucose, were similar. Furthermore, the proliferation of single cells, confined to a plane using a microfluidic device, was followed at the same sampling frequency (every 3 min) using bright-field and fluorescence microscopy and volumes were calculated from cell contours, assuming ellipsoidal geometry. In contrast to our approach they used a super-folding GFP fused to one of the histone 2B loci (HTB2-sfGFP) instead of a bud-neck marker to monitor the cell cycle stage and to discriminate between buds and new born daughter cells. Additionally using a self-developed MATLAB software *Autotrack* for automatized cell segmentation and linage tracking, enabled them to track and analyze thousands of cells.

The cSCGM was fitted to the provided volume trajectories pairs of WT yeast cells, in the same manner as described above. For better comparison with *data 1* the data set was limited to the first cell cycle of new born daughter cells(N= 6079), i.e. cells with a replicative age of 0. Cells, defined as outliers in the data set, were also neglected for further analysis. In contrast to *data 1*, was *t*_budstart_ no fitting parameter but provided by the dataset. For 97 %of the 6079 cells, parameter sets were found for which *χ*^2^ was sufficiently small (Fig. S15). Exemplary volume trajectories, representing data and model fits, are shown in Fig. 5(c) and Fig. S14. Further analysis was restricted to a reduced data set (N=4680), discarding all failed optimization runs, which yield *k*_u/c_ < 1 and *χ*^2^ < 50. The resulting distribution of the fitting parameters and *χ*^2^ is shown in Fig. 5.

**Fig. 5.**
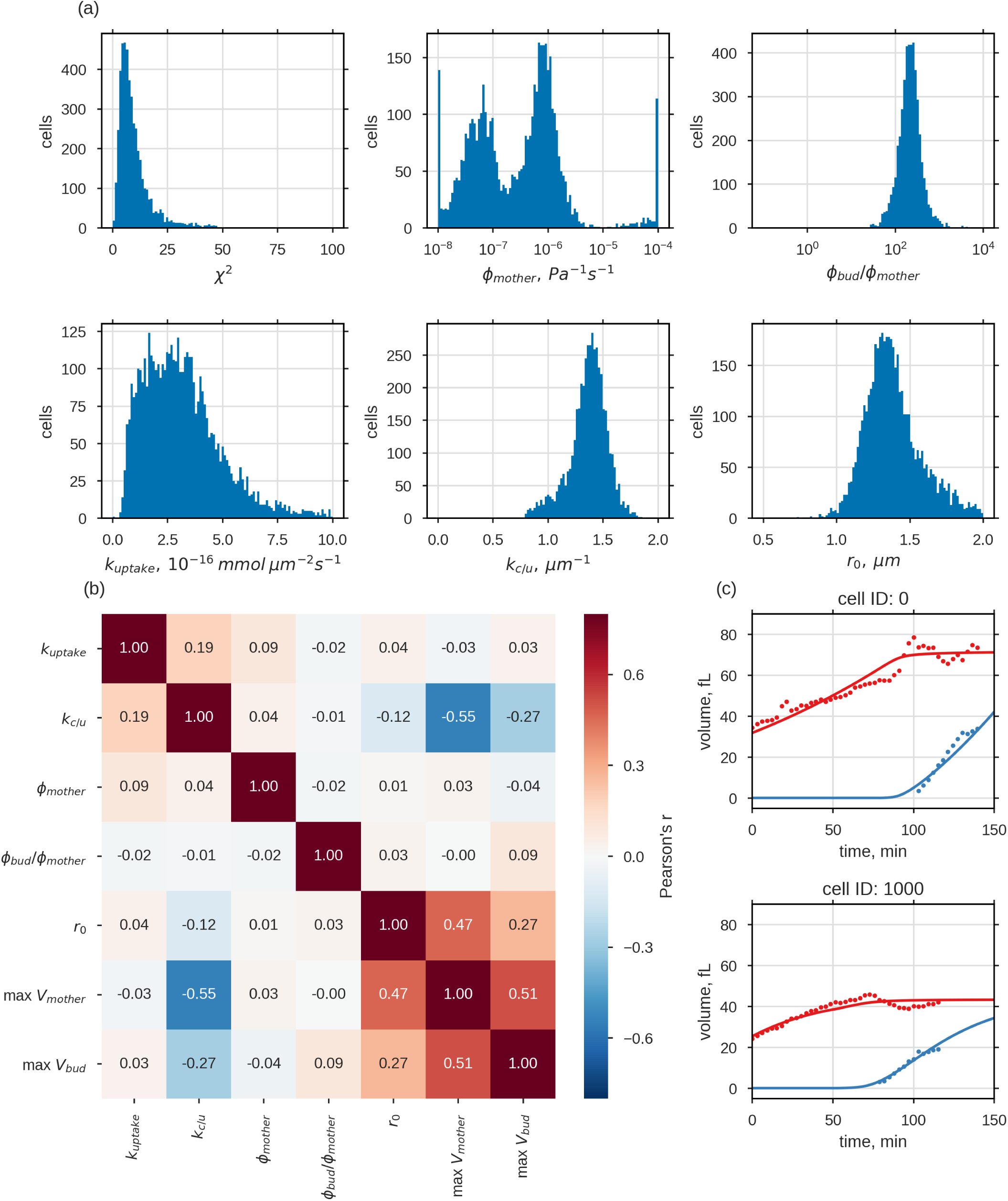
cSCGM fitted reported volume trajectories. **(35)** For 97% of the reported volume data for first-time mothers (N=6079) a parameter set for the cSCGM could be identified. a) Histograms of fitted parameter, excluding parameter sets where *k*_u/c_ < 1 µm or *χ*^2^ > 50 (N=4680, histograms of complete and reduced parameter set shown in Fig. S15). b) Pearson’s correlation of best fitting parameters (*k*_u/c_ < 1 µm, *χ*^2^ > 50) and measured maximal volume of mother and bud, presented as correlogram. max *V*_mother_ and max *V*_bud_ where taken from the data. In c), two exemplary sets of measured and simulated volume trajectories are shown (additional 24 exemplary trajectories are shown in Fig. S14).

#### Parameter distributions

Except for *ϕ*_mother_ the free fit parameter showed a unimodal, though partially skewed, distribution and their medians as well as their interquartile range (IQR) are listed in Tab. 2, together with parameter estimates from *data 1*. Although *t*_budstart_ was not fitted for *data 2*, as it was provided by the data, the parameter is listed in Tab. 2 for comparison.

**Table 2.** Estimated fit parameters of both data sets. *t*_budstart_ of *data 2* was provided by the data. *ϕ*_mother_ showed bimodal distribution and was neglected in this table.

| parameter | unit | factor | median (IQR) |  |
| --- | --- | --- | --- | --- |
|  |  |  | *data 1* (N=21) | *data 2* (N=4680) |
| $k_{\text{uptake}}$ | $\frac{\text{mmol}}{\mu\text{m s}}$ | $10^{-16}$ | 1.4 (1.0–1.9) | 3.0 (1.9–4.2) |
| $k_{\text{c/u}}$ | $\frac{1}{\mu\text{m}}$ | | 1.21 (1.18–1.27) | 1.4 (1.3–1.5) |
| $\phi_{\text{mother}}$ | $\frac{1}{\text{Pa s}}$ | $10^{-4}$ | 5.5 (3.1–6.8) | – |
| $\phi_{\text{bud}}/\phi_{\text{mother}}$ | | $10^2$ | 2.3 (1.3–8.1 $\times 10^3$) | 2.2 (1.5–3.0) |
| $r_0$ | $\mu\text{m}$ | | 1.2 (1.0–1.3) | 1.4 (1.3–1.5) |
| $t_{\text{budstart}}$ | min | $10^1$ | 9.3 (9.3–14.3) | 5.6 (4.1–7.4) |

The extensibility *ϕ*, which reflects the capability of the cell for plastic cell wall expansion and readjustment of the turgor pressure in the SCGM, varied drastically between *data 1* and *data 2*. While *ϕ*_mother_ values obtained from *data 2* followed a clear bimodal-distribution, values obtained from *data 1* showed no indication of a multimodal-distribution. Both maxima, *max*(*ϕ*_mother_|1) = 9.6 × 10^-8^/(Pa s) and *max*(*ϕ*_mother_|2) = 7.8 × 10^-7^/(Pa s) were at least two magnitudes smaller compared to estimates from *data 1* and thereby closer to reported extensibility values for plant cells (∼ 1 × 10^-10^/(Pa s))(17, 36). For each maximum of *ϕ*mother a group of parameter sets was selected, for which *ϕ*mother was close to these maximums, and examined for differences (Fig. S18). Nevertheless no difference in parameter distribution could be observed. Furthermore, screening of volume trajectories for which the estimated *ϕ*_mother_ was either close to *max*(*ϕ*_mother_|1) or *max*(*ϕ*_mother_|2) (Fig. S19, Fig. S20) revealed no common pattern for the two groups. Whether the bimodal distribution of *ϕ*_mother_ results from numerical issues of the optimization algorithm or reflects two distinct population could not yet be finally clarified.

Interstingly, the ratio *ϕ*_bud_*/ϕ*_mother_ (median (IQR)) was comparable between *data 2* (220 (150–300)) and *data 1* (230 (130–8.1 × 10^5^)), underscoring that *ϕ*_bud_ needs to be at least 100 times higher *ϕ*_mother_, regardless the magnitude of *ϕ*, to facilitate bud expansion. Osmolyte uptake rate *k*_uptake_ and ratio *k*_c/u_ estimated from *data 2* were slightly higher than estimates from *data 1*, with *k*_uptake_ = 3.0 (1.9–4.2) 10^-16^ mmol/(μms) and *k*_c/u_ = 1.4 (1.3–1.5) 1/μm, indicating a higher single-cell growth rate and maximal volume for *data 2*.

The initial volume of the new born daughter, i.e. a mother with replicative age 0, is represent by its radius *r*_0_. With *r*_0,1_ = 1.2 (1.0–1.3) μm this radius was slightly smaller for *data 1* than for *data 2*, where *r*_0,2_ = 1.4 (1.3–1.5) μm. There are several possible reasons for the discrepancy between the *r*_0,1_, *r*_0,2_: differences in the genome of investigated strains, experimental setup or the initial growth phase of the population. In contrast to the initial volume, the time until bud emergence *t*_budstart_ was increased for *data 1* compared to *data 2*, with *t*_budstart,1_ = 93 (93–143) min and *t*_budstart,2_ = 56 (41–74) min. Between all fit parameters and the volume maximas of mother and bud (max*V*_mother_ max*V*_bud_), the Pearson’s correlation coefficient was calculated and displayed in a correlogram (Fig. 5(b)). The correlation coefficients can vary by definition between −1 and 1, indicating a negative or positive correlation, whereby a values close to zero represent no correlation. Except for a small correlation (0.19) for *k*_c/u_ and *k*_uptake_ the fit parameter, displayed in the upper left quadrant, were uncorrelated, in contrast to experimental volume measures and *r*_0_ (lower right quadrant), which were all positive correlated. The only correlation between volume measures and fitting parameters, was found for *k*_c/u_. Thereby *k*_c/u_ correlated negatively to all volume measures, though the correlation was much stronger for max*V*_mother_ (−0.55) than for max*V*_bud_ (−0.27) and even less for *r*_0_(−0.12). This can be explained by the cSCGM, since *k*_c/u_ defines the maximal volume of the cell compartment, which is not reached for the bud until after cell separation.

### Comparing active and passive osmotic shock response

Adaptation to changing osmotic conditions is vital for cells. In *S. cerevisiae*, adaptation to high osmolarity is mainly achieved via increased production and retention of the small osmolyte glycerol, coordinated by the high osmolarity glycerol (HOG) pathway, through transcriptional and post-translational regulation of metabolism. We extended the SCGM by a reported description of this HOG cascade, comprising signaling dynamics and the activation of osmolyte production via Hog1 (27). Kinetics in the HOG cascade model were scaled according to the recently measured lower turgor pressure (7). Additionally, we chose *r*_0_ to be at least 1.2*μm* to avoid artificially low Hog1 concentrations. The combined implementation of HOG pathway model and SCGM can be found in the Supplementary Material. Utilizing the augmented model we compared the response of the SCGM to hyperosmotic shock with (Fig. 6, blue lines) and without (Fig. 6, orange lines) active shock response mechanisms mediated by HOG signaling.

**Fig. 6.**
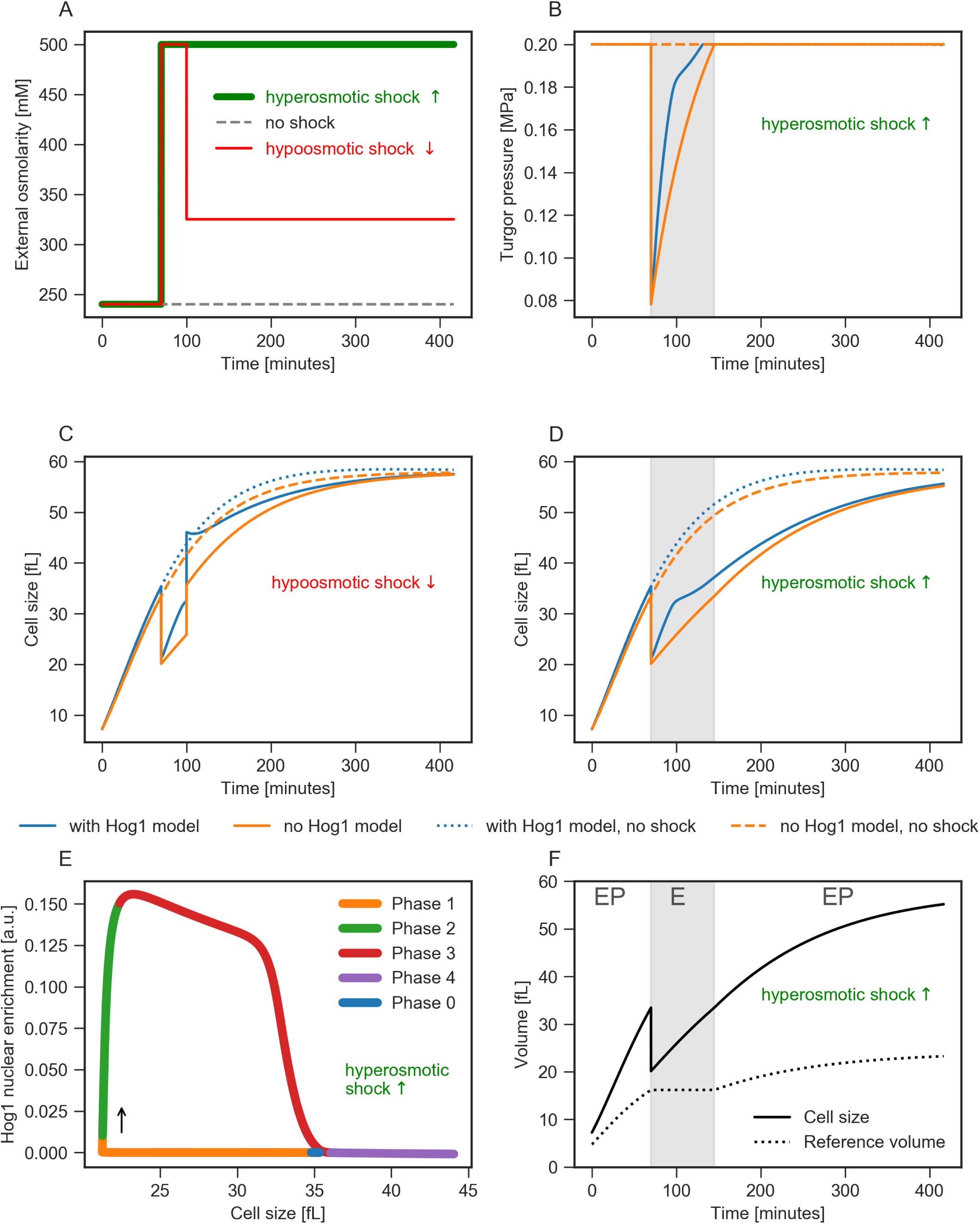
Description is on the next page. Growing single cell exposed to osmotic shocks. Simulation of our SCGM with a HOG response model reproduces Hog1 signaling behavior in growing yeast cells. We compared SCGM dynamics upon hyper- and hypo-osmotic shock with and without active response mechanism. **A** Shock scenarios: no shock (black dashed line), single hyperosmotic shock (external osmolarity raised to 500 mM at 70 minutes, green line), and combination of hyper- and hypo-osmotic shock (external osmolarity is raised to 500 mM at 70 minutes and reduced to 325 mM at 100 minutes, red line). (**B-D**) Shock response (**B, D** - only hyperosmotic, **C** hyper- and hypo-osmotic) with (blue lines) and without (orange lines) active HOG response. **B** shows turgor pressure, which drops and re-adapts with and without signaling, but faster with signaling. **C, D** show cell size, which also adapts faster to stress in case of active HOG response. **E** Short- and long-term response of the model with HOG response to the same hyperosmotic shock as in **B, D** showing Hog1 nuclear enrichment versus cell size trajectory as four distinct phases of response behavior after shock (indicated as phase zero) as proposed by (26): (1) passive but fast shrinkage and turgor loss, (2) activation of Hog1 signaling, (3) restoration of cell size and turgor pressure, and finally (4) resuming cellular growth. Our model reflects these phases and combines the fast timescale in (1) (volume loss within a few seconds) and steady growth either without a shock or due to adaption after shock as in the last phase. **F** Comparison of cell size to reference volume (shock as in **B** and **D**). The reference volume reflects the distinct cell wall mechanic regimes of our model: elasto-plastic behavior (EP) during growth, ideal elastic behavior (E, grey area) in response to hyperosmotic shock, and growth resumption (EP) after restoring former size and turgor pressure (see turgor trajectory in **B**).

When unperturbed (in Fig. 6, dashed lines), both SCGM and SCGM with HOG establish steady states for internal osmolarity and turgor pressure while sharing comparable growth rates. However, if we simulate a hyperosmotic shock as depicted in Fig. 6A, the models show an identically fast decrease in volume and turgor and a drastic increase in *c*_*i*_ (see Supplementary Material for the *c*_*i*_ trajectory), but the following trajectories diverge. The active response of the HOG cascade leads to production and accumulation of glycerol, which contributes to fast reestablishment of former turgor pressure and cell size (blue line). Without HOG cascade, cells still manage to adapt volume and turgor, but much slower. A systematic analysis of the effects of strength and timing of osmotic shock is provided in Supplementary Material Fig. S24 and Fig. S23.

The SCGM with HOG can reproduce previous measurements of growing yeast cells that undergo a hyperosmotic shock. In (26), cellular volume versus Hog1 localization as a shock response was measured over time for individual yeast cells, resulting in a phase plot that is reproduced by our model simulations (Fig. 6E). In particular, the characteristics throughout the four different phases that are reflected correctly by our approach, validate that not only the HOG response model or our volume model are functional by themselves but, more importantly, that interplay of both models is captured properly. Further, we tested whether the model also copes with hypoosmotic (low external osmolarity) shock (Fig. 6C). Again model integrity is restored following a first hyperosmotic shock and, thus, regains osmolarity and volume after a second, now hypoosmotic, shock. Obviously, hypoosmotic shock results in a short overshoot of cell size (Fig. 6C), as previously reported in (37). Hyperosmotic shock exclusively leads to elastic deformation, while hypoosmotic shock leads to negligibly small plastic deformation. Both types of osmotic shock induce elastic behavior of the cell wall, hence the modeled cell would not grow effectively through step-like osmotic changes of the environment. The reference volume (Fig. 6F and Supplementary Material for detailed definition) quantifies growth as it integrates plastic deformation of the cell wall under normal conditions reflecting a elasto-plastic behavior (EP) (Fig. 6F, white area). In contrast, a hyperosmotic shock induces immediate cell shrinkage and stops growth, as indicated by the reference volume. The reference volume remains constant, since the elasto-plastic regime is left and the model behaves ideal-elastically (E) (Fig. 6F, grey area).

## Discussion

Knowledge of single-cell growth dynamics is crucial for the understanding of comprehensive biological systems, particularly for model systems such as *S. cerevisiae*. Here, we introduced and justified a volume model of yeast by integrating different theoretical concepts of cell mechanics, integrity and osmolyte dynamics in combination with experimental data on single cell growth and cell wall elasticity. The model applies to several scenarios, where cell size variation plays a role: from growth of an individual bud connected to the mother cell, the case of a two-bud growing mutant of yeast, and the direct and long-term volume response after hypertonic osmotic shock.

In previous studies, two modes of single-cell growth were considered for yeast, linear and exponential (38–41). Based on our model we proposed a more complex picture of the specific growth pattern of a single yeast cell (a comparative plot of the three modes of single-cell growth can be seen in Fig. 1B). We derived an exact analytic solution for the radius trajectory depending on time *r*(*t*) for a single cell growing undisturbed. Prospectively, this solution may support future work as it relates relevant parameters, which originate from formerly distinct concepts such as cell wall mechanics, water homeostasis and osmoregulation in the form of a single expression describing the growth of an individual cell overall. Such an expression allows us to deduce properties of the described system in a closed form. Specifically, we exemplified such a deduction by calculating the limes of the radius for long time spans *r*_*t*→∞_. The approach revealed that the maximal cell size is determined by the ratio *k*_u/c_ of osmolyte or nutrient uptake and consumption, but is independent of mechanical cell wall properties, such as Young’s modulus or extensibility. The SCGM works fully without a connection to the cell cycle machinery comprising the action of cyclins, CDKs and CK1. Nevertheless, the coupled SCGM is not independent of cell cycle, since the characteristic length of first cell cycle is integrated via *t*_budstart_. Parameterizing the SCGM with measured volume trajectories we found that this characteristic cell cycle length varied significantly between individual cells.

Yeast cells proliferate by budding, a process in which a larger mother and a smaller bud grow simultaneously but with different growth rates. To describe this coordinated growth, two instances of the SCGM with different initial volumes were coupled by allowing water and osmolyte fluxes between the instances. For simplicity, we neglected the specific bud localization at the mother, which would include the complex polarization process (42, 43). This coupled SCGM can be interpreted as an augmented model compared to a previous description of interconnected soap bubbles or balloons (30, 44). However, those models lack the exchange of matter with their environment which is crucial for biological applications. In the coupled SCGM we allowed fluxes inbetween the cavities and between cavities and medium such that our model operates near equilibrium instead of being at equilibrium. Plastic and elastic cell wall expansion depends on the lateral stresses, which scales with radius and acting pressure. To allow expansion of the smaller of the two volumes, the cell wall of that volume must be either more elastic (*E*) or more extendable (*ϕ*).

Utilizing atomic force microscopy, we could reject the hypothesis that elasticity is the distinguishing cell wall property between mother and bud. The measured local Young’s modulus at the bud was not lower but similar or even slightly higher compared to the mother. Therefore, we inferred from the SCGM that *ϕ* must be significantly higher at the bud to facilitate bud expansion. Fitting the coupled SCGM to the growth data suggests that *ϕ*_mother_ is three orders of magnitude higher than reported values for plants (45, 46) and *ϕ*_bud_ needs to be at least two magnitudes higher than at mother cell wall. Furthermore, we have shown that significantly increased *ϕ*_bud_ compared to *ϕ*_mother_ can explain the observed higher growth rates of buds (31). The higher extensibility of the bud cell wall might be caused by incorporation of new more expandable material, which maturates over time or by subsequent alterations of the already formed cell wall. Plant enzymes are reported to alter mechanical cell wall properties (47, 48), thereby influencing growth and cell shape (49) and reports also suggest that cell wall plasticity in fungi is controlled by hydrolytic enzymes, like chitinases or glucanases (5, 50–52).

Fitting the coupled model to single-cell volume trajectories from two completely independent data sets enabled us to estimate crucial growth parameters, such as the global osmolyte uptake rate *k*_uptake_ and the ratio *k*_u/c_. Both parameters appeared to be very sensitive, as *k*_uptake_ had the strongest impact on individual growth rate and *k*_u/c_ on the maximum cell volume. From the comparison of both data sets two main observations can be stated: First, cell expansion of *S. cerevisiae*, under non-limiting conditions, requires a total osmolyte uptake rate of 1.0 to 4.0 mmol/(μm^2^ s) and a consumption rate which is 1.2 to 1.5 μm^-1^ times higher. Second, the extensibility of the bud cell wall has to be at least 100 higher than the mother’s, to facilitate bud expansion.

Intriguingly, the estimated osmolyte uptake rates *k*_uptake_ were only two to five times higher than the average glucose uptake rate per surface area under aerobic conditions reported by (53) (see supplementary note 5). Under these conditions glucose would therefore account for 30 % to 60 % of the osmolytes taken up by the cell, revealing a new angle on the impact of glucose onto cellular growth: So far the impact of glucose onto cell growth based exclusively on the fact that glucose is a main nutrient source, providing chemical energy and building blocks for macromolecules, and hence can be regarded as fuel of the cell. While this study indicates, that additionally to those aspects, the massive glucose import could drive water influx and hence contribute significantly to cell expansion.

In general, the cSCGM described very well the volume expansion of different single-cell trajectories, which have been measured in microfluidic growth experiments. Using the cSCGM, we could also recapitulate earlier experimental data presented by (41) and the resulting relations between division time and ratio of volumes at division and birth (Fig. S10), despite different interpretation for the growth dynamics of single cells.

By integrating the osmotic stress response, specifically the HOG signaling pathway and metabolic adaptation from (27), in the SCGM, we could compare the passive volume adaptation with the active response arising from the HOG pathway. This model was able to capture the different time scales ranging from steady growth to the drastic shrinkage of cell size followed by an osmotic shock. To our knowledge, this is the first time that the joint-dynamics of Hog1 enrichment versus volume following a hyperosmotic shock (26) could be reproduced and explained by a mathematical model. Additionally, our model can describe the response to hypoosmotic shock similarly to previous reports (37). The simulations illustrate the functionality of the coupled model over a wide range of cell sizes. However, validity of the combined model is limited in cases of tiny initial volume (e.g. 1 fL), since the Zi-model assumes fixed numbers of Hog1 molecules. During growth, increasing volume results in dilution of all species within the HOG model and, hence, would lead to non-physiologically small concentrations for tiny initial volumes. To overcome such limiting effects, production rates for all species of the HOG model need to be introduced and parameterized to counteract the dilution process. Instead, we chose a simplistic implementation of coupling both models, in order to preserve their original properties.

In this study, we focused on the single-cell growth of newly budded yeast cells and described their volume development. Extending the cSCGM to cover several cell cycles with several consecutive budding events and test it against experimental data might help to understand the interplay of morphogenesis and aging of *S. cerevisiae*. An other step would be the combination of the SCGM with models focusing on cell cycle regulation or polarization into larger models to investigate the interplay between regulatory networks and growth. Mathematical models for growth dynamics at the population level are often based on greatly simplified single cell volume models (54, 55). The presented SCGM might help to improve such models and, hence, increase our understanding of population growth dynamics.

## Methods and Materials

### Yeast strains

For the microfluidic growth analysis we used a *Saccharomyces cerevisiae* strain based on BY4742 (MAT*α*his3Δ1 leu2Δ0 lys2Δ0 ura3Δ0) (56), in which the bud neck marker Cdc10 has been genomically labeled with mKate2. The sequence of mKate2 was cloned into the plasmid pUG72 (Euroscarf) and Cdc10 was labeled using PCR-based homologous recombination. The necessary Ura3 marker cassette has been removed using the Cre-loxP recombination system (57). In the nano-indentation experiments we used the strain BY4741 (56).

### Yeast cell culture

For the microfluidic growth experiments, cells were grown at 30 °C overnight in synthetic medium (SD; 0.17% yeast nitrogen base without amino acids, 0.5% ammonium sulfate, 2% glucose, 55 mg/L adenine, 55 mg/L l-tyrosine, 55 mg/L uracil, 20 mg/L l-arginine, 10 mg/L lhistidine, 60 mg/L l-isoleucine, 60 mg/L l-leucine, 40 mg/L ysine, 10 mg/L l-methionine, 60 mg/L phenylalanine, 50 mg/L l-threonine and 40 mg/L l-tryptophan). Medium osmolarity was measured with an osmometer (gonotec, Berlin, Germany). Before the experiment, 500 *μ*L of overnight culture were diluted in 5 ml SD medium and grown for 2.5 hours at 30 °C.

### Microfluidic growth experiments

We used the CellASIC ONIX microfluidic platform (Merck Millipore, Darmstadt, Germany) with the haploid yeast plates (Y04C) for growth analysis. Plates were primed with SD medium and cells were loaded as described in the ONIX yeast protocol. Flow control pressure was set to 2 psi. Cells were observed using a Visitron Visiscope inverted spinning disc laser confocal microscope (Visitron, Puchheim, Germany). The temperature of the microfluidic plate was controlled to 30 °C using a temperature control chamber (OL IX73/83 cellVivo, PeCon GmbH, Erbach, Germany). We used a 150X oil immersion objective (Olympus UPlanSApo 150X/1.47, Oil, TIRM) and the Photometrics Evolve 512 EMCCD camera (Photometrics, Tuscon, USA). Fluorescent illumination was provided by a 561 nm diode laser and a multibad dichroic filter (405/488/559/635 nm) together with a 600/50 nm emission filter for detection. A brightfield image was taken every 3 minutes, while a fluorescent image was only taken every 12 minutes, to minimize fluorescent exposure of cells. Each image was acquired as a z-stack (6 z-positions, 0.5 *μm* distance).

### Image analysis

For each z-stack, the sharpest plane was determined using the ImageJ plugin “Find focused slices” by Quingzong Tseng. Brigthfield images were segmented and cells were tracked with the CellProfiler (58) plugin Cellstar (34). Tracking was manually checked, and mis-tracked cells were removed from the analysis. For each cell, bud appearance time was manually defined as the time when a bud was first visible in the brightfield image. The lineage was manually determined using the fluorescent images of the bud neck marker Cdc10-mKate2. Cell area was converted to volume by assuming a spherical cell shape.

### Cell wall nano-indentation

In brief, log-phase yeast cells were mechanically trapped in porous polycarbonate membrane, with pore size of 5 *μm* (59). The cell-containing membrane was subsequently immobilized on the bottom of a liquid chamber and probed with an AFM (Nanowizard III, JPK, Germany). All AFM measurements were conducted in liquid solution of synthetic media using MLCT-E cantilever (Bruker). Prior to all experiments, the spring constant was determined via thermal noise method. Multiparametric imaging of cells was done using the provided QI-Mode, whereby maximal applied force was set to 1 nN, corresponding to an indentation depth of ∼35 nm. For further analysis we used JPK data processing software the Python packages numpy, scipy (60) or pandas (61). For a more detailed description see (7).

## ACKNOWLEDGEMENTS

This work was supported by the Deutsche Forschungsgemeinschaft (DFG) through Collaborative Research Center (SFB740, to EK) and by project “Osmotic stress regulation” (UH 279/1-1, to JU).

