## Supplementary Material for "Osmolyte homeostasis controls single-cell growth rate and maximum cell size of *Saccharomyces cerevisiae*"

### Supplementary Note 1: Derivation of the differential equation for turgor pressure

Here, we deduce an ordinary differential equation (ODE) for turgor pressure in a walled cell by using the mechanical formalism that was introduced above (see Fig. 1). The thin shell relation provides a conversion from turgor pressure $\Pi_t$ to the cell wall in-plane stress $\sigma$. For a spherical geometry it reads:

$$\sigma = \frac{\Pi_t r}{2d}, (r \gg d) \tag{1}$$

and holds for a cellular radius $r$ much bigger than cell wall thickness $d$. Both principle circumferential strains $\varepsilon_c = \varepsilon_\theta = \varepsilon_\phi$ are given by:

$$\mathrm{d}\varepsilon_c = \frac{\mathrm{d}r}{r} \tag{2}$$

The mechanical response of the cell wall to the acting turgor pressure can be modeled by two principle mechanical elements, a Hookean and a Bingham-Norten element connected in series. While the stress remains equal for all elements, $\sigma = \sigma_{Hook} = \sigma_{Bingham}$ the strain is distributed among the elements

$$\mathrm{d}\varepsilon = \frac{\mathrm{d}r}{r} = \frac{\mathrm{d}r_{Hook}}{r} + \frac{\mathrm{d}r_{Bingham}}{r}. \tag{3}$$

Resulting in two constitutive relationships for stress and strain or stress and strain-rate for the elastic or plastic deformation, respectively.

$$\varepsilon_{Hook} = \frac{1-\nu}{E}\frac{\Pi_t r}{2d}, \tag{4}$$

$$\dot{\varepsilon}_{Bingham} = \frac{\phi r}{2d} f_m(\Pi_t, \Pi_c), \quad \text{where} \begin{cases} \Pi_t, & \text{if } \Pi_t > \Pi_c \\ 0, & \text{else} \end{cases} \tag{5}$$

Here $\phi$ is the plastic extensibility and $\Pi_c$ is the critical turgor pressure above which the plastic expansion occurs. If the acting turgor pressure is below $\Pi_c$ only elastic expansion will take place. Hereinafter, a dot above a variable will represent the time derivative of that variable $\dot{x} = \frac{\mathrm{d}x}{\mathrm{d}t}$.
Considering Eq. 5, Eq. 3 and the time derivative of Eq. 4

$$\dot{\varepsilon}_{Hook} = \frac{1-\nu}{E2d}(\dot{\Pi}_t r + \Pi_t \dot{r}), \tag{6}$$

the strain rate with respect to cell radius can be written as

$$\frac{\dot{r}}{r} = \frac{1-\nu}{2Ed}(\dot{\Pi}_t r + \Pi_t \dot{r}) + \frac{\phi r}{2d} f_m(\Pi_t, \Pi_c). \tag{7}$$

From here we can finally deduct the differential equation for the turgor pressure

$$\dot{\Pi}_t = \frac{2Ed}{1-\nu}\frac{\dot{r}}{r^2} - \Pi_t\frac{\dot{r}}{r} - \frac{E\phi}{(1-\nu)} f_m(\Pi_t, \Pi_c). \tag{8}$$

For further analysis we defined a reference radius $R_{ref}$, which is constraint to be equal to the relaxed cellular radius when $\Pi_t < \Pi_{crit}$, which allows us to follow the plastic deformation over time, e.g. the growth. Using Eq. 5 the strain rate of $R_{ref}$ is given by

$$\frac{\dot{R}_{ref}}{R_{ref}} = \frac{\phi r}{2d} f_m(\Pi_t, \Pi_c). \tag{9}$$

Collecting this term in Eq. 8 leads to the ODE that was implemented

$$\dot{\Pi}_t = \frac{2Ed}{1-\nu}\left(\frac{\dot{r}}{r^2} - \frac{1}{r}\frac{\dot{R}_{ref}}{R_{ref}}\right) - \Pi_t\frac{\dot{r}}{r}. \tag{10}$$

Furthermore, the Kedem-Katchalsky equation provides the osmotic changes of the radius for spherical cells when rewritten to

$$\dot{r} = -L_p(\Pi_t + \Pi_e - \Pi_i). \tag{11}$$

This way the ODE for turgor pressure is a combination of two concepts, water homeostasis of the cell and cell wall mechanics.

### Supplementary Note 2: Integration of ideal elastic turgor ODE results in a turgor-radius relation that is similar to the Merritt-Weinhaus equation

As mentioned in the main article we can show that our turgor description, specifically the ideal elastic version of Eq. 8 shares similarities with the Merritt-Weinhaus equation (44). Our turgor description Eq. 8 reduces to an ideal elastic version of the turgor pressure ODE (e.g. during a hyperosmotic shock), thus yielding a function for $\Pi_t(r)$ depending on cellular radius, describing the pressure-to-size relation of an ideal elastic thin shell.

In order to compare both descriptions of turgor we need to integrate our turgor ODE first. To solve the ideal-elastic turgor ODE, which is Eq. 8 when setting the term that contains the $max$-function to zero

$$\dot{\Pi}_t = \frac{2Ed}{1-\nu}\frac{1}{r^2}\frac{\mathrm{d}r}{\mathrm{d}t} - \frac{\Pi_t}{r}\frac{\mathrm{d}r}{\mathrm{d}t},$$

one can rewrite this as a differential of $\Pi_t(r(t),t)$ according to radius dependency

$$\frac{\mathrm{d}\Pi_t}{\mathrm{d}r} = \frac{2Ed}{1-\nu}\frac{1}{r^2} - \frac{\Pi_t}{r}.$$

Considering the condition for the turgor to be zero at $r_{ref}$, $\Pi_t(r_{ref}) = 0$ yields:

$$\Pi_t(r) = \frac{2Ed}{1-\nu}\left(\frac{ln(r)}{r} - \frac{ln(r_{ref})}{r}\right),$$

in fact $r_{ref}$ is the reference radius, which can be integrated by simulating the complete set of ODEs as shown in the main article. For our simulations we estimated the initial value of the reference radius based its relation to the initial turgor pressure and initial cellular radius by rewriting the latter equation: $r^0_{ref} = f(\Pi^0_t, r^0)$.

### Supplementary Note 3: Integration of Kedem-Katchalsky equation

In this section, we present an analytical solution of our entire ODE model focusing on the volume variation while turgor and osmotic pressure are kept in steady-state (e.g. a non-perturbed growing cell).

As a summary, the ODE system contains these three equations for internal osmolarity, turgor and volume:

$$\frac{\mathrm{d}c_i}{\mathrm{d}t} = k_{uptake}\frac{G}{V} - k_{consumption}\frac{V}{V} - c_i\frac{1}{V}\frac{\mathrm{d}V}{\mathrm{d}t} \qquad \textbf{(12)}$$

$\frac{\mathrm{d}\Pi_t}{\mathrm{d}t}$ : (see Equation 8 from above.)

$$\frac{\mathrm{d}r}{\mathrm{d}t} = -L_p(\Pi_t + \Pi_e - RTc_i) \qquad \textbf{(13)}$$

In order to solve this system we will try to collapse the system into a single ODE of this kind:

$$F(\dot{r}, c_i^{SS}, \Pi_t^{SS}, t) = 0, \qquad \textbf{(14)}$$

where this differential equation depends on radius $r$, time-derivate of the radius $\dot{r}$, time $t$ and quasi-steady-states for internal osmolarity $c_i^{SS}$ and turgor pressure $\Pi_t^{SS}$ and thus meeting our constrains for an not perturbed growing single cell.

Hence, we calculate the quasi-steady-state of the internal osmolarity $\dot{c}_i = 0$ yielding

$$c_i^{SS} = \frac{\Pi_t + \Pi_e}{2RT} + \sqrt{\frac{(\Pi_t + \Pi_e)^2}{(2RT)^2} + \frac{k_{\text{uptake}} - \frac{k_{\text{consumption}}}{3}r(t)}{RTL_p}}.$$

Thus, the quasi-steady-state of the internal osmolarity depends on the turgor $\Pi_t$, the external osmotic pressure $\Pi_e$, the osmolyte uptakte rate constant $k_{\text{uptake}}$, the osmolyte consumption rate $k_{\text{consumption}}$, the hydrostatic permeability of the cell membrane $L_p$ (that either have constant values or are in their steady-states) and on the cellular radius $r(t)$ which is time dependent. Thus, for a growing cell size $c_i$ has only a quasi-steady-state, $\dot{c}_i \approx 0$. This is an important property and becomes apparent in our solution below.

In particular, using this $c_i^{SS}$ inside the Kedem-Katchalsky equation:

$$\frac{\mathrm{d}r}{\mathrm{d}t} = -L_p\left(\Pi_t + \Pi_e - \left(\frac{\Pi_t + \Pi_e}{2} + \sqrt{\frac{(\Pi_t + \Pi_e)^2}{4} + \frac{(k_{upt.} - \frac{k_{cons.}}{3}r)RT}{L_p}}\right)\right)$$

$$\frac{\mathrm{d}r}{\mathrm{d}t} = -L_p\frac{\Pi_t+\Pi_e}{2} + \sqrt{\frac{L_p^2(\Pi_t+\Pi_e)^2}{4} + k_{upt.}L_pRT - \frac{k_{cons.}L_pRT}{3}r},$$

Considering the turgor pressure $\Pi_t$, numerical simulations of our ODE model suggest that turgor is also in steady-state ($\Pi_t^{SS}$) if the cell grows without any osmotic stress. Specifically, we assume the steady-state value of the turgor to be close to the critical turgor pressure, $\Pi_t^{SS} \approx \Pi_{tcrit.} = 0.2\ MPa$.
In the latter ODE for $r_t$ we can lump the constants:

$$\alpha := -L_p\frac{\Pi_t+\Pi_e}{2}$$
$$\beta := \frac{L_p^2(\Pi_t+\Pi_e)^2}{4} + k_{upt.}L_pRT$$
$$\gamma := \frac{k_{cons.}L_pRT}{3},$$

where we can identify an ODE that can be solved:

$$\frac{\mathrm{d}r}{\mathrm{d}t} = \alpha + \sqrt{\beta-\gamma r}$$
$$\int \frac{\mathrm{d}r}{\alpha+\sqrt{\beta-\gamma r}} = \int \mathrm{d}t$$

An integral would be:

$$\int \frac{\mathrm{d}r}{\alpha+\sqrt{\beta-\gamma r}} = \frac{2(\alpha\log(\sqrt{\beta-\gamma r}+\alpha)-\sqrt{\beta-\gamma r})}{\gamma} + const. = F(r) + const.$$

Solving the integral for r by including lower and upper limits:

$$F(r) = (t-t_0) + F(r_0) \quad \textbf{(15)}$$

and rearrangement towards $r$ yields a time-dependent function for the radius:

$$r(t) = \frac{1}{\gamma}\left(-\alpha^2+\beta-2\alpha^2 W\left(-\frac{1}{\alpha}e^{\frac{\gamma(F(r_0)+(t-t_0))}{2\alpha}-1}\right) - \alpha^2 W^2\left(-\frac{1}{\alpha}e^{\frac{\gamma(F(r_0)+(t-t_0))}{2\alpha}-1}\right)\right), \quad \textbf{(16)}$$

where $W$ is the Lambert-$W$ function and we summarized

$$F(r_0) = \frac{2(\alpha\log(\sqrt{\beta-\gamma r_0}+\alpha)-\sqrt{\beta-\gamma r_0})}{\gamma},$$

with the initial radius $r_0$ at the initial time $t_0$.
Interestingly, the exponential inside the two occurrences of $W$ functions in our solution converges against zero for large $t$ and subsequently the two terms containing the $W$ function vanish for large t:

$$r_{final} := \lim_{t\to\infty} r(t) = \frac{-\alpha^2+\beta}{\gamma} = 3\frac{k_{upt.}}{k_{cons.}},$$

which describes the final radius of the cell.

### Supplementary Note 4: Deduction of a simplified version of our radius equation.

In the previous section we deduced a function for the development of the radius of a cell over time by solving our ODE-system analytically. The exact solution, as written in Eq. 16, is rather complex. It would be useful to have a simplified relation at hand that adequately describes a growing single cell. Here, we deduce such a simplified relation. During deduction we checked the validity of each of our simplifications within the range of physiological reasonable radii and times by evaluating our radius equation Eq. 16 for the proposed set of parameters (Tab. 1) and using this as a reference as shown in Fig. S7.
Starting with a first order Taylor series approximation for the Lambert-W function as the inner exponential is close to zero for our parameter set and times $t \geq 0$. Note, the Taylor approximation of Lambert-W function $W(\cdot)$ around zero results the unity function:

$$r(t) \approx \frac{1}{\gamma}\left(-\alpha^2+\beta-2\alpha^2\left(-\frac{1}{\alpha}e^{\frac{\gamma(F(r_0)+(t-t_0))}{2\alpha}-1}\right)-\alpha^2\left(-\frac{1}{\alpha}e^{\frac{\gamma(F(r_0)+(t-t_0))}{2\alpha}-1}\right)^2\right). \quad \textbf{(17)}$$

Furthermore, the exponential squared (term in Eq. 17) is much smaller than the exponential (un-squared in the middle) and also smaller than the other terms in the latter equation. Hence, omitting this term on the far right of the latter equation yields:

$$Eq.\ 17 \approx \frac{1}{\gamma}\left(-\alpha^2+\beta-2\alpha^2\left(-\frac{1}{\alpha}e^{\frac{\gamma(F(r_0)+(t-t_0))}{2\alpha}-1}\right)\right). \quad \textbf{(18)}$$

Writing out the $F(r_0)=\frac{2(\alpha\log(\sqrt{\beta-\gamma r_0}+\alpha)-\sqrt{\beta-\gamma r_0})}{\gamma}$ in detail and collecting our previously defined $r_{final} := \frac{-\alpha^2+\beta}{\gamma}$:

$$Eq.\ 18 = r_{final} + \frac{2\alpha}{\gamma}\left(e^{(\log(\sqrt{\beta-\gamma r_0}+\alpha)-\frac{1}{\alpha}\sqrt{\beta-\gamma r_0}}e^{\frac{\gamma(t-t_0)}{2\alpha}}e^{-1}\right), \quad \textbf{(19)}$$

$$= r_{final} + \frac{2\alpha}{\gamma}\left((\sqrt{\beta-\gamma r_0}+\alpha)e^{-\frac{1}{\alpha}\sqrt{\beta-\gamma r_0}}e^{\frac{\gamma(t-t_0)}{2\alpha}}e^{-1}\right). \quad \textbf{(20)}$$

where the major bracket can be separated into a time-dependent part and a time-independent term:

$$= r_{final} + \frac{2\alpha}{\gamma}\left((\sqrt{\beta-\gamma r_0}+\alpha)e^{-\frac{1}{\alpha}\sqrt{\beta-\gamma r_0}-1}\right)e^{\frac{\gamma(t-t_0)}{2\alpha}}. \quad \textbf{(21)}$$

The time-independent exponential in the latter equation can be approximated by this relation: $2e^{x-1} \approx x+1$ for values of x close to one, which is true for our underlying parameter set ($-\frac{1}{\alpha}\sqrt{\beta-\gamma r_0} \approx 1$). (The alternative use of a Taylor approximation (e.g. $2e^{x-1} \approx 2x$) would be a more precise approximation but is rather simplifying the latter relation and therefore was rejected.)

$$Eq.\ 21 \approx r_{final} + \frac{\alpha}{\gamma}\left((\sqrt{\beta-\gamma r_0}+\alpha)(1-\frac{1}{\alpha}\sqrt{\beta-\gamma r_0})e^{\frac{\gamma(t-t_0)}{2\alpha}}\right). \quad \textbf{(22)}$$

$$= r_{final} + \left(\frac{-\beta+\alpha^2}{\gamma}+r_0\right)e^{\frac{\gamma(t-t_0)}{2\alpha}}. \quad \textbf{(23)}$$

Again collecting our definition for $r_{final} := \frac{-\alpha^2+\beta}{\gamma}$ yields:

$$= r_{final} - \left(r_{final}-r_0\right)e^{\frac{\gamma(t-t_0)}{2\alpha}}, \quad \textbf{(24)}$$

which is our proposed equation describing the growth of a single cell as a trajectory of radius:

$$\tilde{r}(t) = r_{final} - \left(r_{final}-r_0\right)e^{\frac{\gamma(t-t_0)}{2\alpha}}. \quad \textbf{(25)}$$

In agreement to our exact solution, for large times $t$, the $\tilde{r}$ tends towards $r_{final}$, which describes the final radius of the cell and $r_0$ is the initial radius of the cell. Writing out the lumped constants $\alpha, \beta, \gamma$ and $r_{final}$ explicitly by applying their definitions from above yields:

$$\tilde{r}(t) = r_{final} - \left(r_{final}-r_0\right)e^{\frac{-k_{uptake}RT}{r_{final}(\Pi_t+\Pi_e)}(t-t_0)}. \quad \textbf{(26)}$$

Note, for the exponent one can rewrite the definition of $r_{final}$ like this: $k_{consumption}/3 = k_{uptake}/r_{final}$.

### Supplementary Note 5: Estimation of glucose uptake rate per surface area

According to (53) the mean glucose uptake rate of two replicates of *Saccharomyces cerevisiae* is $k_{glc} = 1.06\ mmol\ (g\ dry\ wt)^{-1}\ h^{-1}$. When additionally considering the reported values from Klis et al. (62) for dry weight per cell ($m_{cell} = 16.5\ pg\ dry\ wt/cell$) and average surface ($A = 60\mu m^2$), we get an estimation of the glucose uptake per surface area:

$$k_{\text{glc,up}} = \frac{k_{glc}\ m_{cell}}{A}$$
$$= \frac{1.06\ mmol\ 16.5\ 10^{-12} g\ dry\ wt}{3.6\ 10^3\ s\ 60\ \mu m^2\ g\ dry\ wt}$$
$$\approx 0.8\ 10^{-16}\ mmol\ \mu m^{-2}\ s^{-1}$$

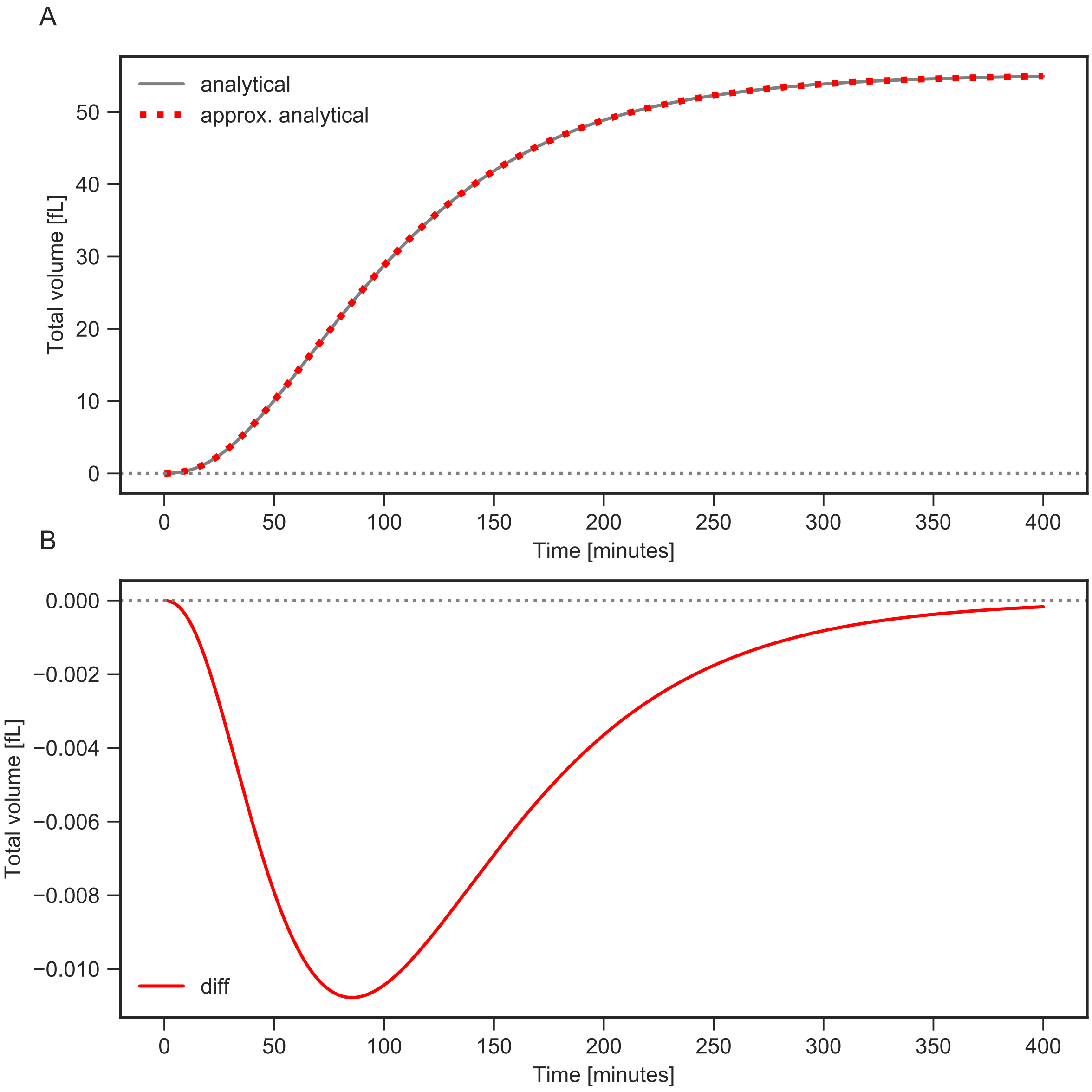


**Fig. 7. Exact analytical solution versus simplified description of cell size.** A: Cell size over time (solid line) by $V(r(t)) = 4/3\pi r(t)^3$ with r(t) described by our analytical solution Eq. 16 versus our simplified radius description $\tilde{r}(t)$ (dotted line). B: Estimating error by calculating the difference between both graphs in A. Obviously, the initial state and the limes state (for large time t) are equal for both solutions, whilst errors appear mostly during the transition (extrema at $\approx 90\,\mathrm{min}$). Overall, the resulting errors of total volume due to approximation/simplification of r(t) are much smaller than the absolute values of total volume (compare the magnitudes on the y-axis between A and B).

### Supplementary Note 6: SCGM with HOG: Integration of our single cell growth model with a HOG cascade model

To include cell signaling, specifically the HOG1 signaling cascade we augmented our single cell growth model with the HOG response model reported by (27). Specifically, we had to adjust the kinetics in the HOG cascade model. Those were scaled according to the recently measured lower turgor pressure (7). Additionally, we chose $r_0$ to be at least $1.2\mu m$ to avoid artificially low Hog1 concentrations. The combined implementation of HOG pathway model and SCGM can be found in our code repository (https://github.com/tbphu/volume_model).

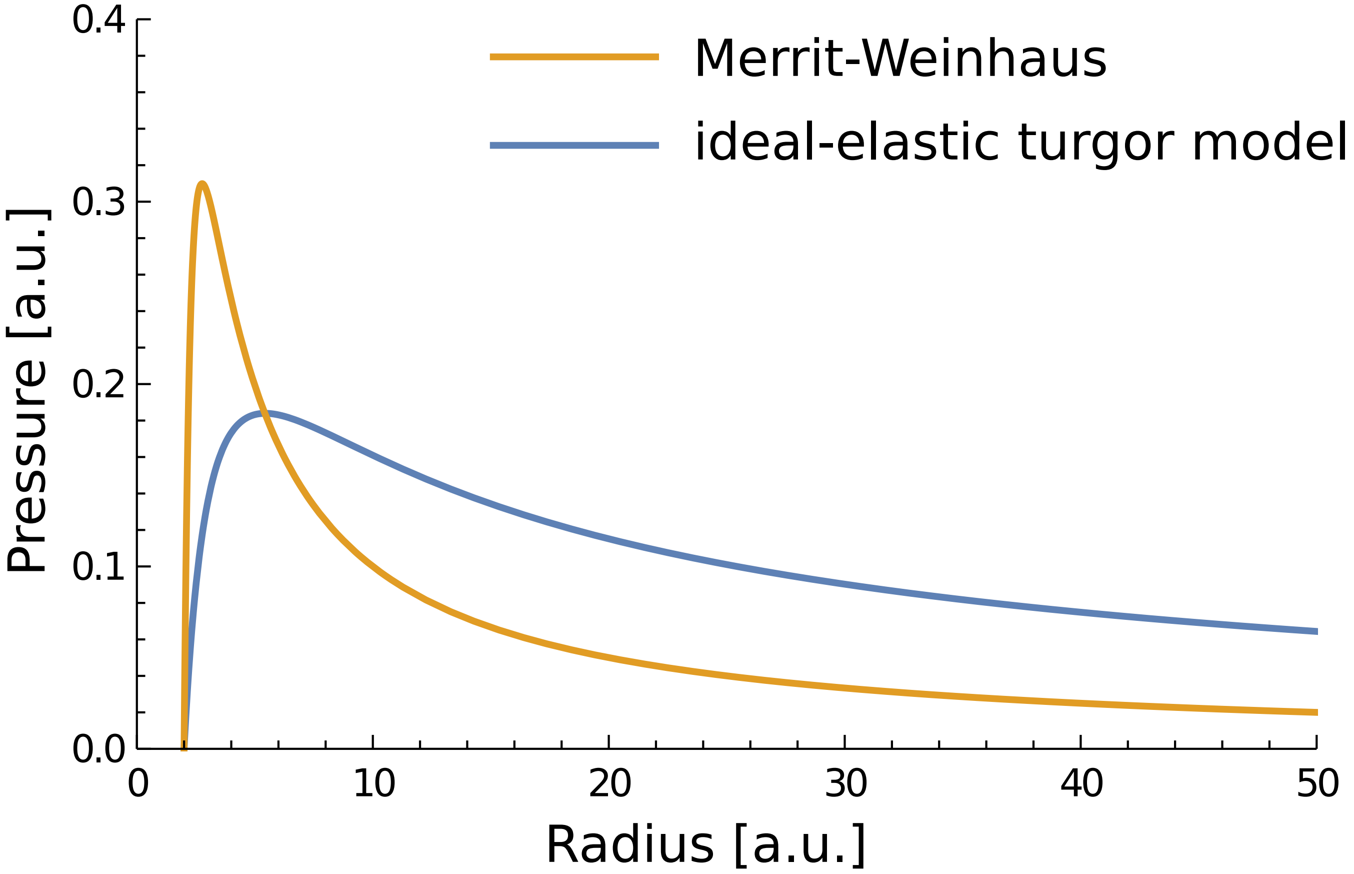


**Fig. 8. Comparison of the Merritt-Weinhaus equation and our ideal-elastic turgor model of a single yeast cell.** The Merritt-Weinhaus equation $\Pi(r) = k(\frac{1}{r} - \frac{r_0^6}{r^7})$ is parameterized with $k = 1$ and $r_0 = 2$. Our model shares the same generic parameters: $\frac{2Ed}{1-\nu} = 1$ and $r_0 = 2$ to ensure comparability in this specific plot. Both models show a similar overall behavior.

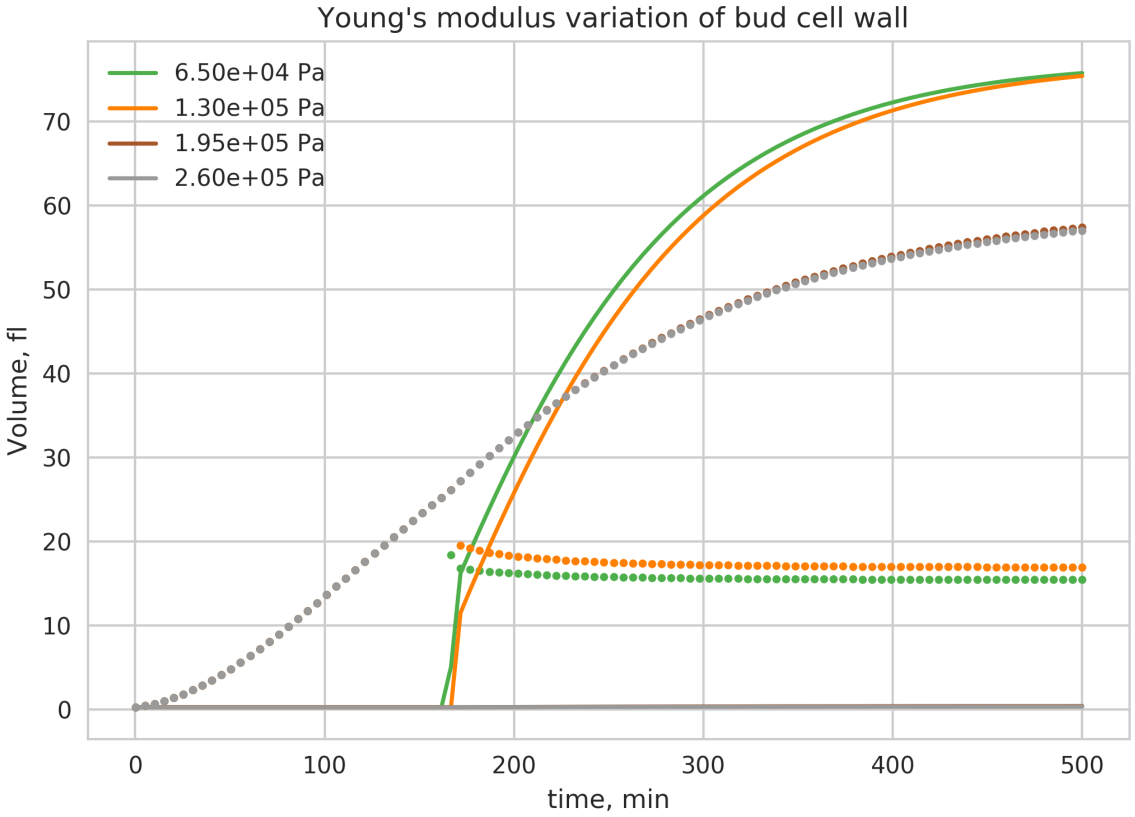


**Fig. 9. Strongly reduced Young's modulus could result in bud growth.** Growth of mother and bud according to the model with identical extensibilities in both compartments, solid line represents bud volume and dotted line represents mother volume. Only small values (< 195 kPa) Young's modulus of the bud cell wall, in comparison to the mother, the bud starts to grow.

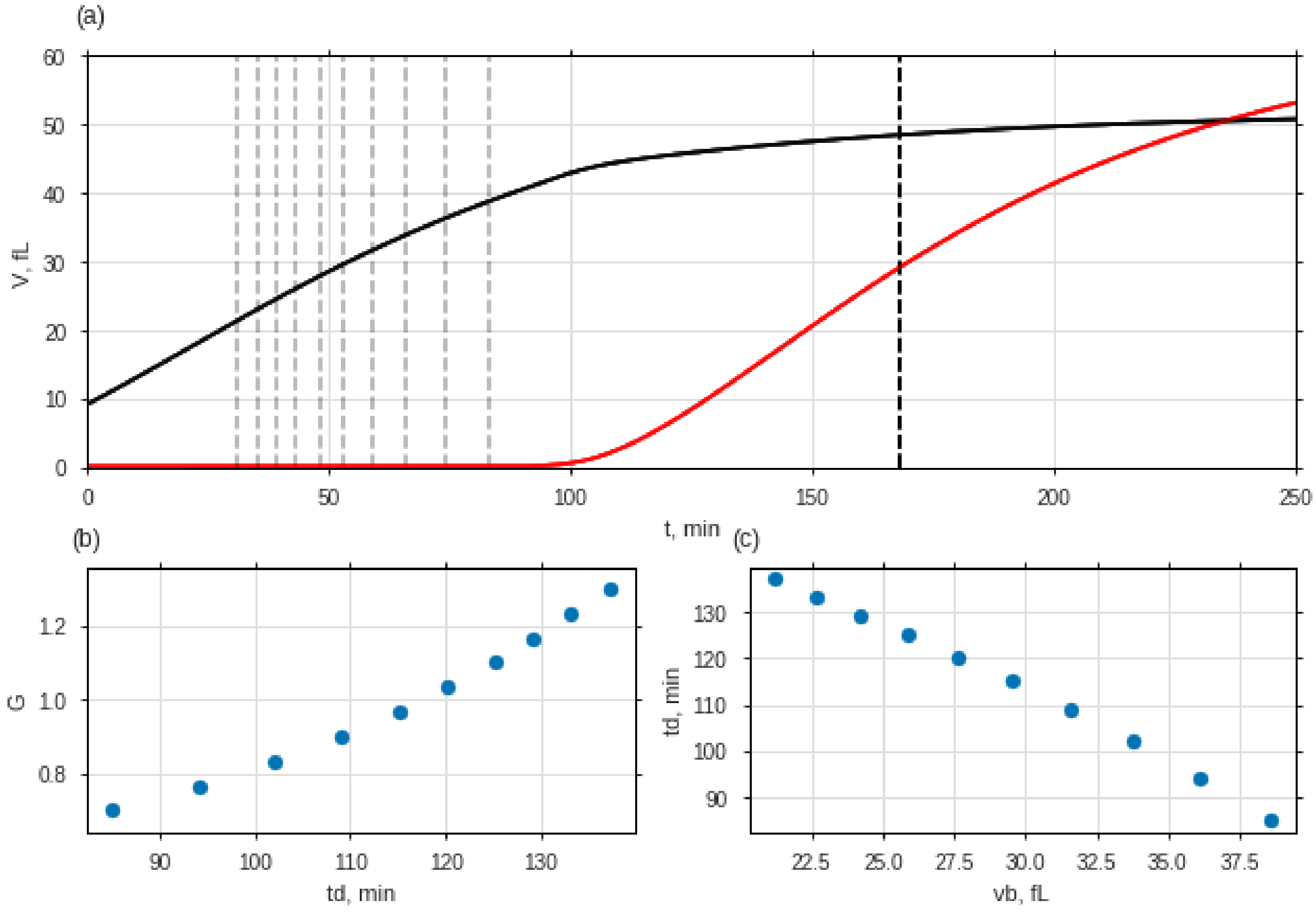


**Fig. 10.** The coupled SCGM can recapitulate the observation form (41).(a) Volume trajectories of mother (black) and bud (bud) calculated with the coupled SCGM are shown. In (b) the logarithm of the volume ratio between the whole cell $vd$ and the previous bud $vb$ is shown. ($G = log(vd/vb)$) Panel (c) shows the estimated division time $td$ vs the initial bud size $vb$. Additional to earlier made assumptions we used a ratio of 0.6 between mother and bud to define the division time $td$ and sampled for different $G$-values between 0.7 and 1.3.

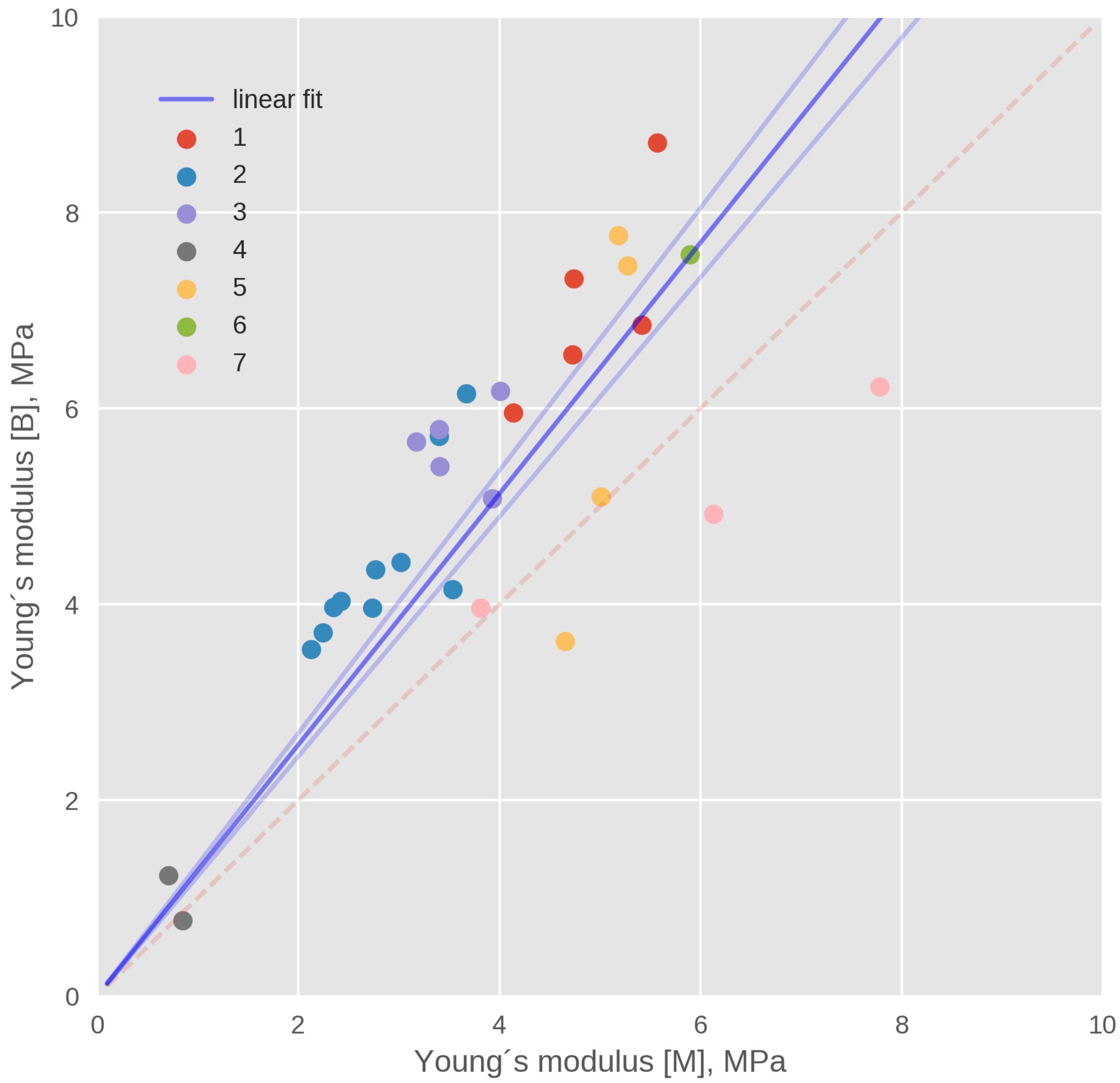


**Fig. 11. Young's modulus of the bud with respect to the mother for seven different cells.** The blue line represents the linear fit with no offset, while the light blue line correspond to upper and lower standard deviation. The calculated slope was 1.28 +/- 0.06. Colors correspond to individual cells, measured at different points in time.

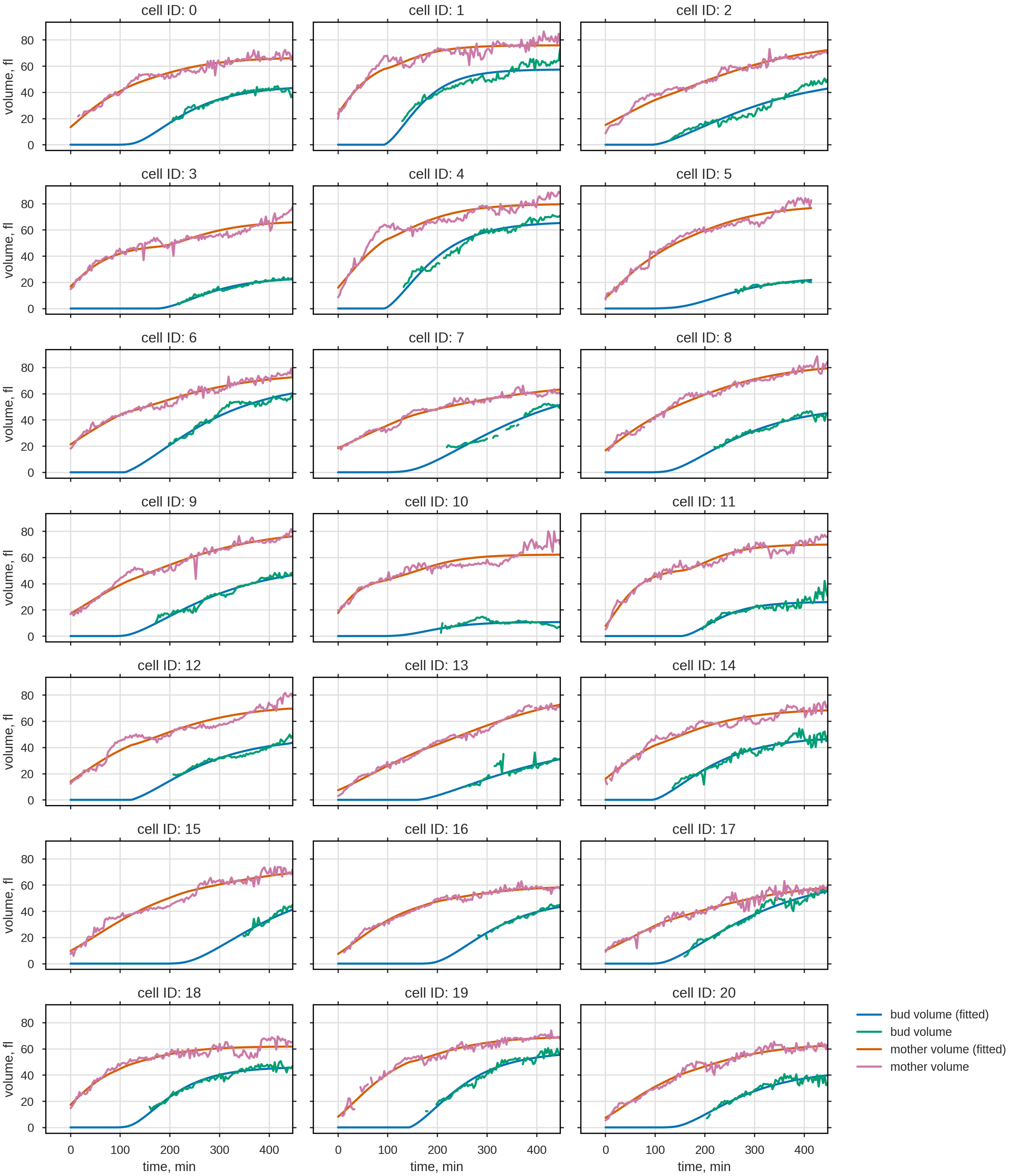


**Fig. 12. Volume trajectories of mother and bud of all 21 analyzed cells and the corresponding volume trajectories from the fitted coupled SCGM.** The volume evolution of 21 individual yeast cells was recorded using bright-field microscopy and microfluidics in combination with semi-automatized image recognition. The segmentation algorithm demands an minimal size, hence the first appearance of the bud was delayed.

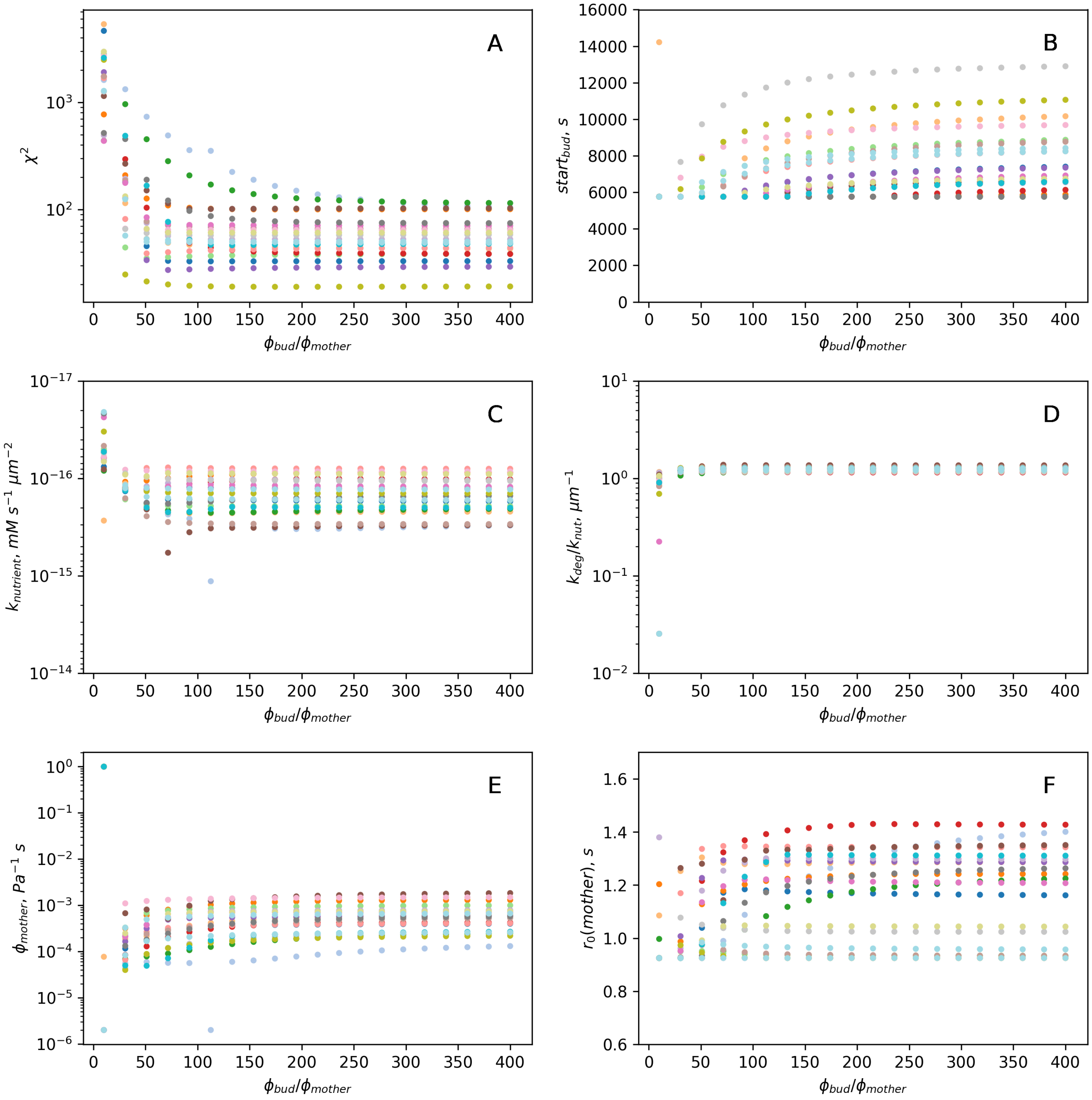


**Fig. 13. Profile likelihood and sensitivity of $\phi_{\mathbf{bud}}/\phi_{\mathbf{mother}}$.** **A** From the profile likelihood of the expansion rate ratios $\phi_{bud}/\phi_{mother}$ we could infer that $\phi_{bud}$ needs to be at least 100 times higher than $\phi_{mother}$, although an upper bound could not be determined. Sensitivity of the fitted free parameters **B-F** to $\phi_{bud}/\phi_{mother}$ shows that only the starting conditions (osmotic $r_0$, $t_{budstart}$,) respond to an variation while the key parameters $k_{nut}$, $k_{deg}/k_{nut}$, and $\phi_{mother}$ remain stable for small $\chi^2$ values. Colors correspond to individual cells.

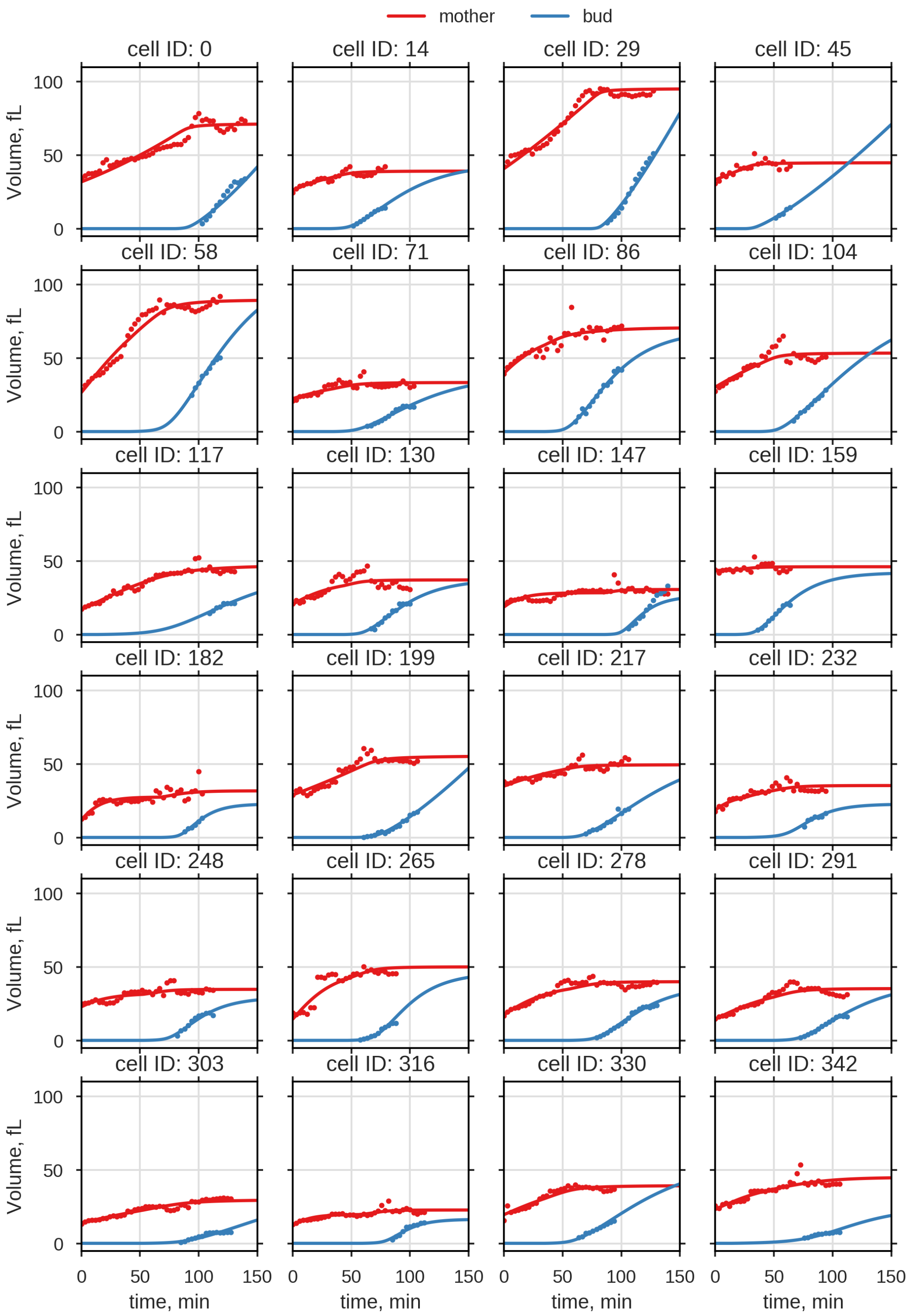


**Fig. 14.** 24 analyzed volume trajectories, out of 5880 provided by (35), and corresponding fits of the cSCGM.

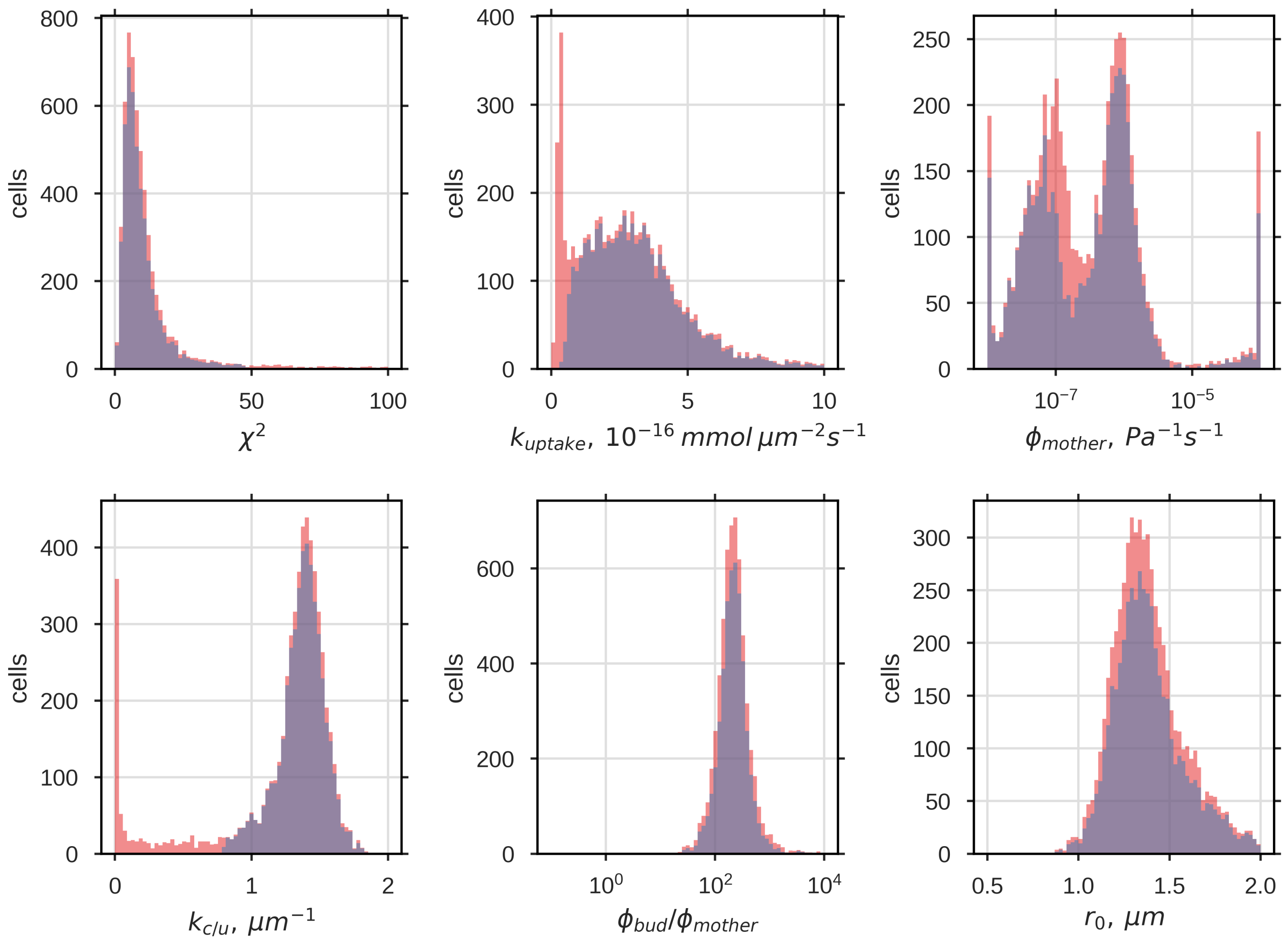


**Fig. 15.** Distributions of the estimated model parameters and $\chi^2$. In red distribution of all parameter set for all fitted cells (N=5880) and in blue the reduced parameter set for which $k_{\mathrm{u/c}} > 0.8\,\mu\mathrm{m}$ and $\chi^2 < 50$ (N=4680). First peak of $k_{\mathrm{uptake}}$ results solely from fits where $k_{\mathrm{u/c}}$ is below 0.8 µm. Discarding sets with high $\chi^2$ and low $k_{\mathrm{u/c}}$ shifts the first maximum of $\phi_{\mathrm{mother}}$ to lower values while the second remains unaltered.

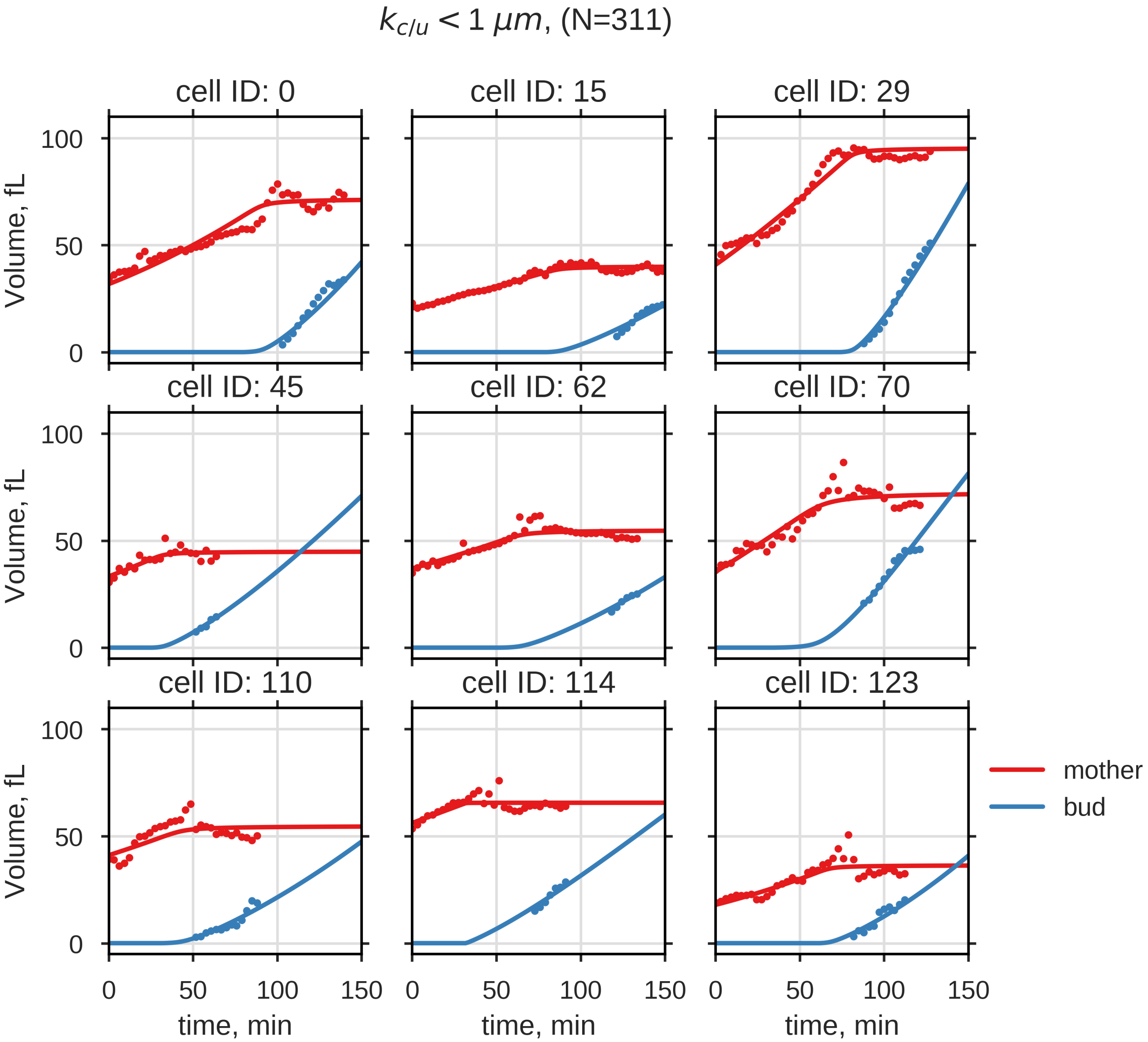


**Fig. 16.** Nine exemplary bud volume trajectories with corresponding fits, where $k_{\mathrm{u/c}} < 1\,\mu\mathrm{m}$. Simulated bud volume shows no decreased growth rate at the end of the cell cycle.

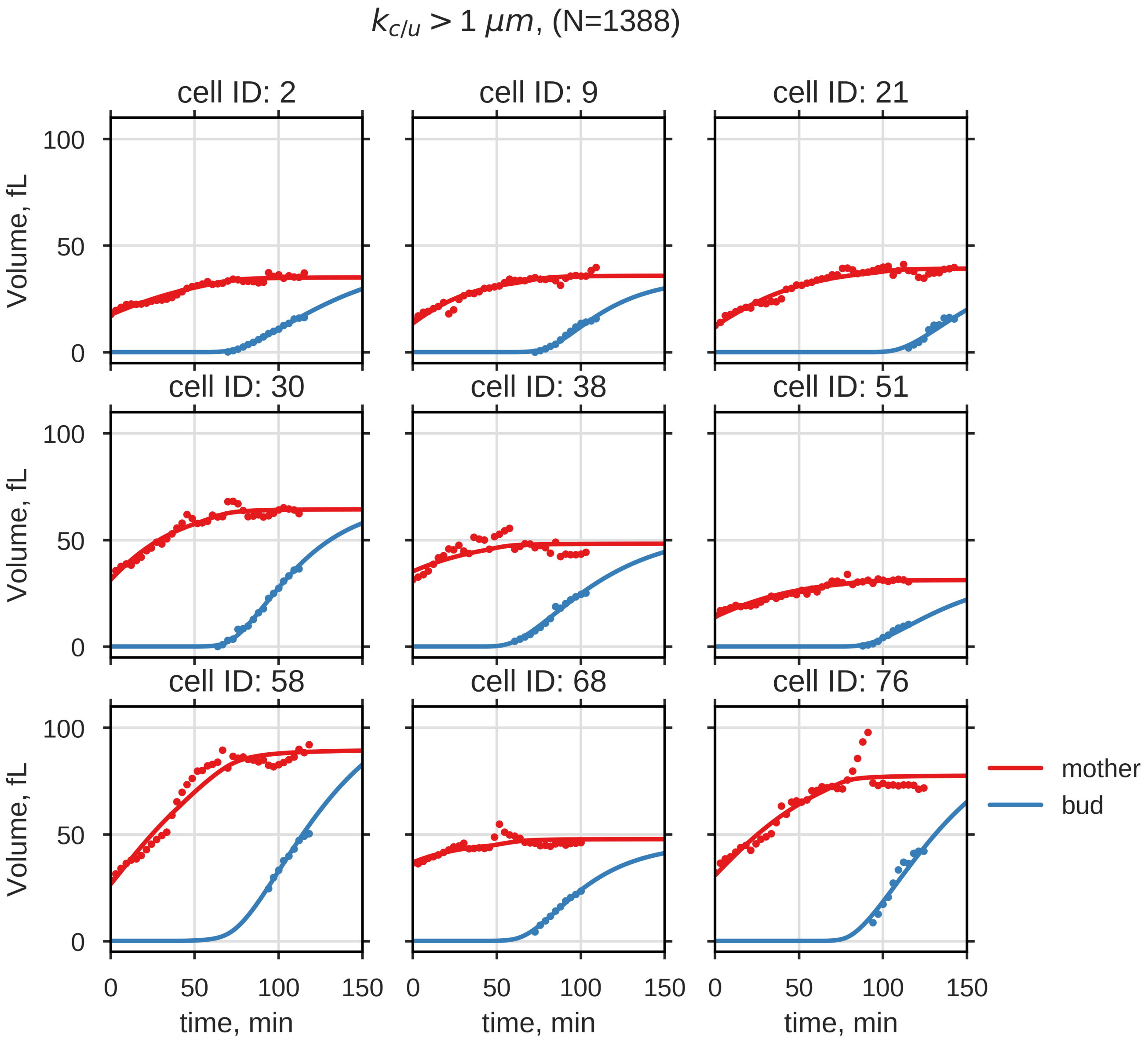


**Fig. 17.** Nine exemplary bud volume trajectories with corresponding fits, where $k_{\mathrm{u/c}} > 1\,\mu\mathrm{m}$

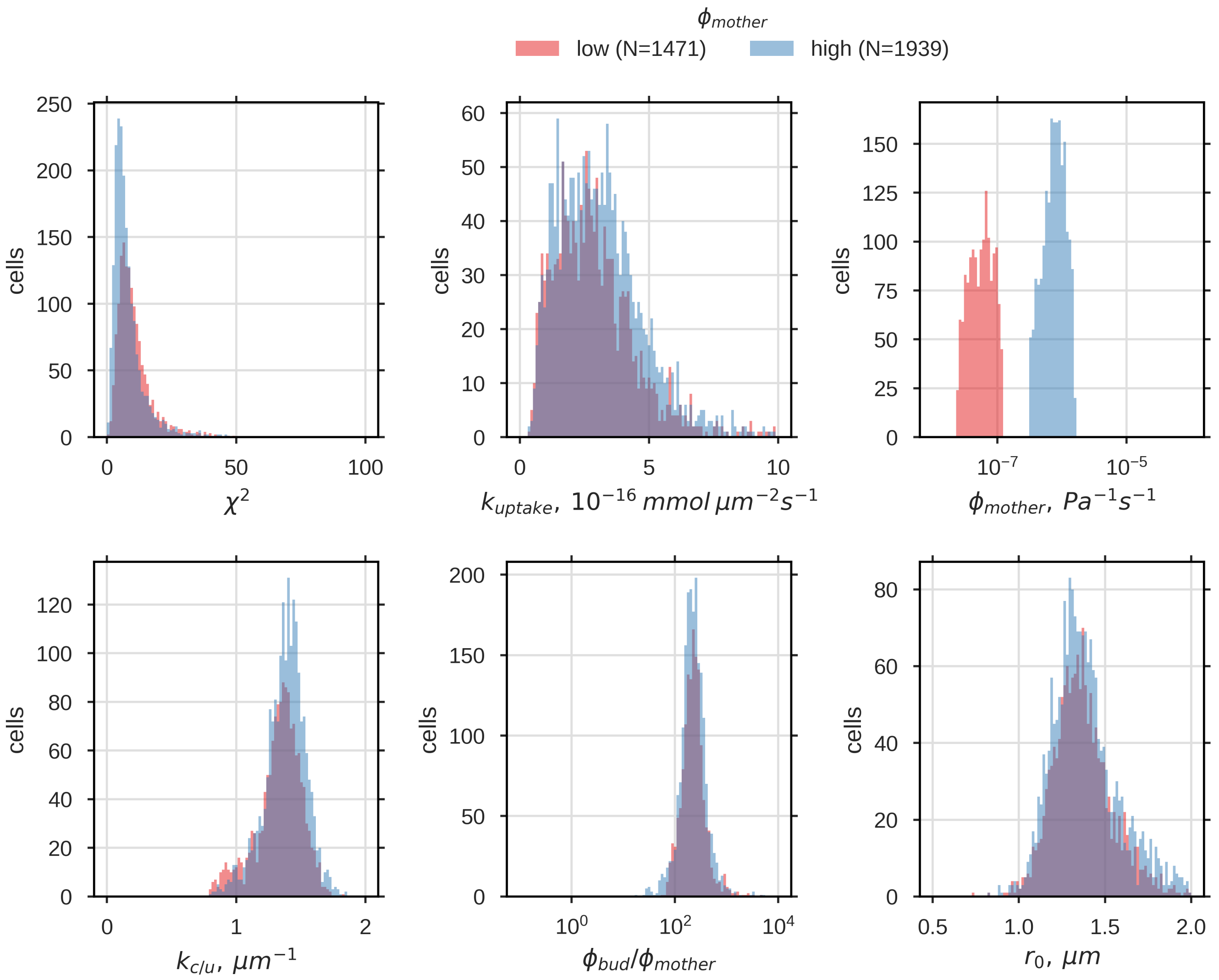


**Fig. 18.** Distributions of the estimated model parameters and $\chi^2$ for $\phi_{\text{mother}} \approx \mathbf{max}(\phi_{\text{mother}}|\mathbf{1})$ or $\mathbf{max}(\phi_{\text{mother}}|\mathbf{2})$, red or blue.

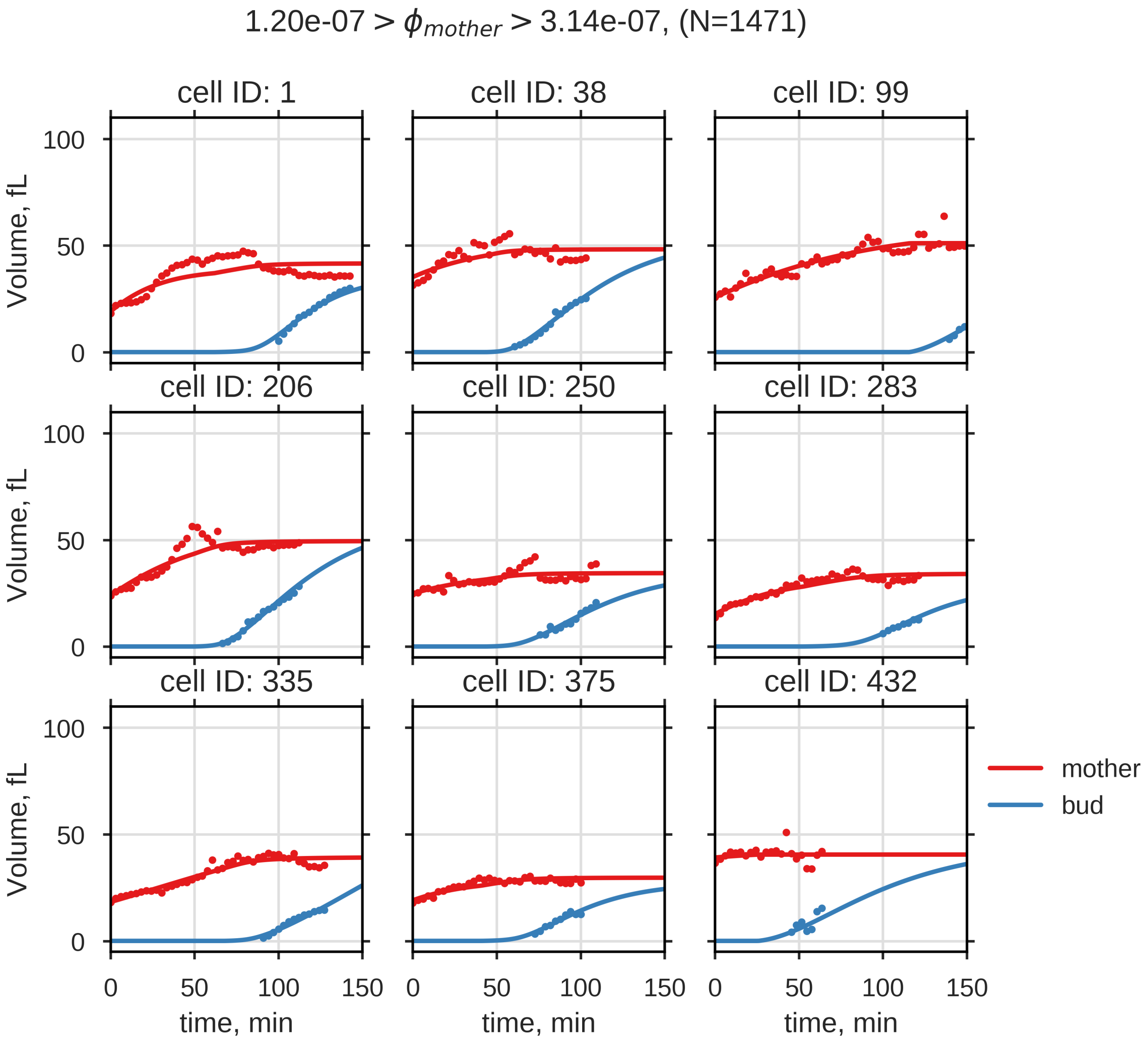


**Fig. 19.** Nine exemplary bud volume trajectories with corresponding fits, where $\phi_{\text{mother}}$ is close to $\phi^{max}_{\text{mother}}(1)$

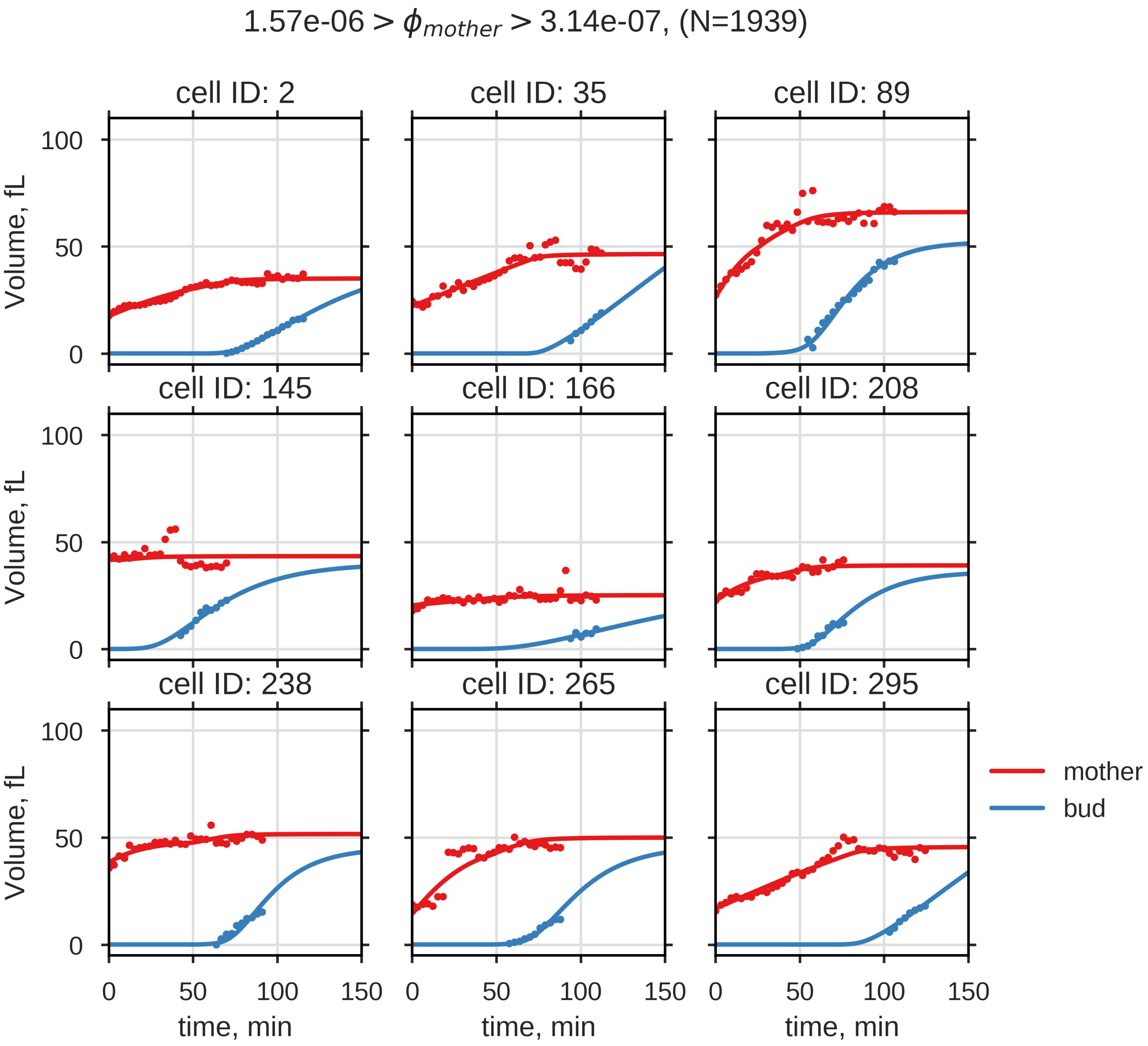


**Fig. 20.** Nine exemplary bud volume trajectories with corresponding fits, where $\phi_{\text{mother}}$ is close to $\phi^{max}_{\text{mother}}(2)$

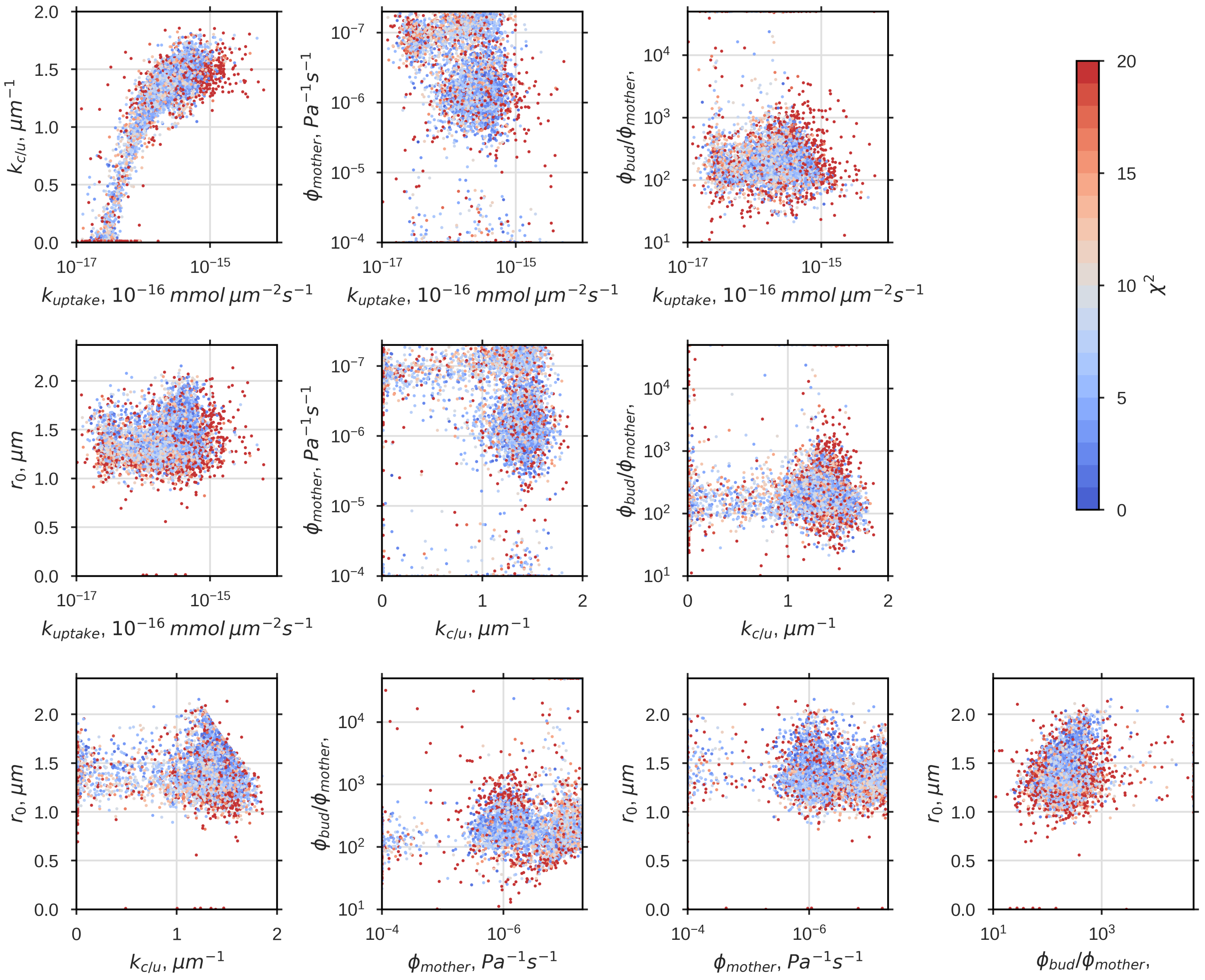


**Fig. 21.** Correlation between fitted parameters. Color represents $\chi^2$

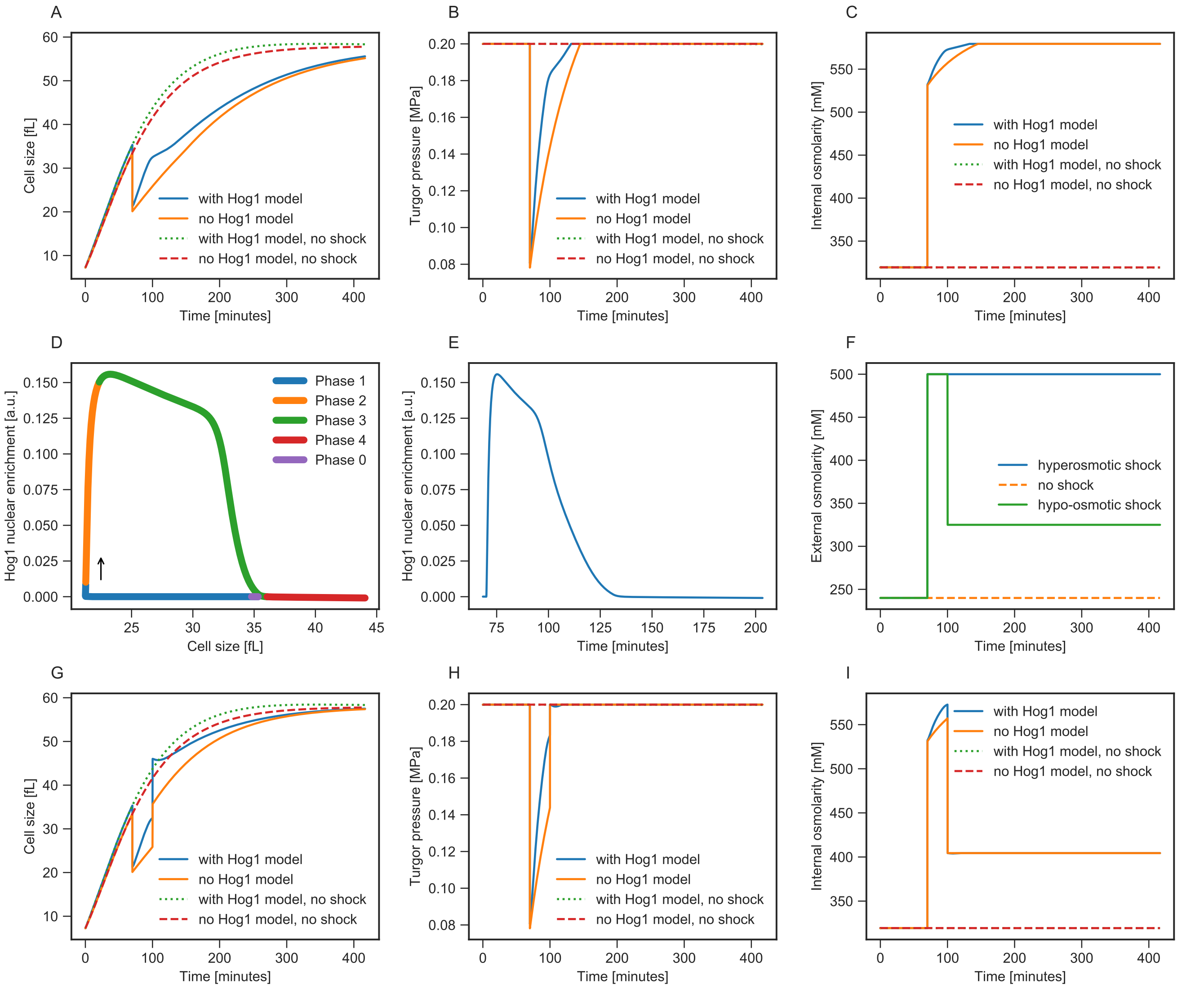


**Fig. 22. Growing single cell exposed to osmotic shocks (Equivalent to the respective figure in the main article but with additional plots, e.g. internal osmolarity)** Top row, A-C: cell size, turgor pressure and internal osmolarity for a single hyperosmotic shock (at 70 minutes, external osmolarity is raised to 500 mM). D and E: Hog1 nuclear enrichment (definition according to Muzzey 2009) as response to the hyperosmotic shock. F: External osmolarity shock profiles for hyperosmotic (blue line, applies to A-E) and hyper- and hypoosmotic shock (green line, applies to G-I). For the hyperosmotic shock: the external osmolarity is raised to 500 mM at 70 minutes. For the hypoosmotic shock: the external osmolarity is dropped back to 325 mM at 100 minutes, green line in E). Bottom row, G-I: cell size, turgor pressure and internal osmolarity for a hyperosmotic shock followed by a hypoosmotic shock.

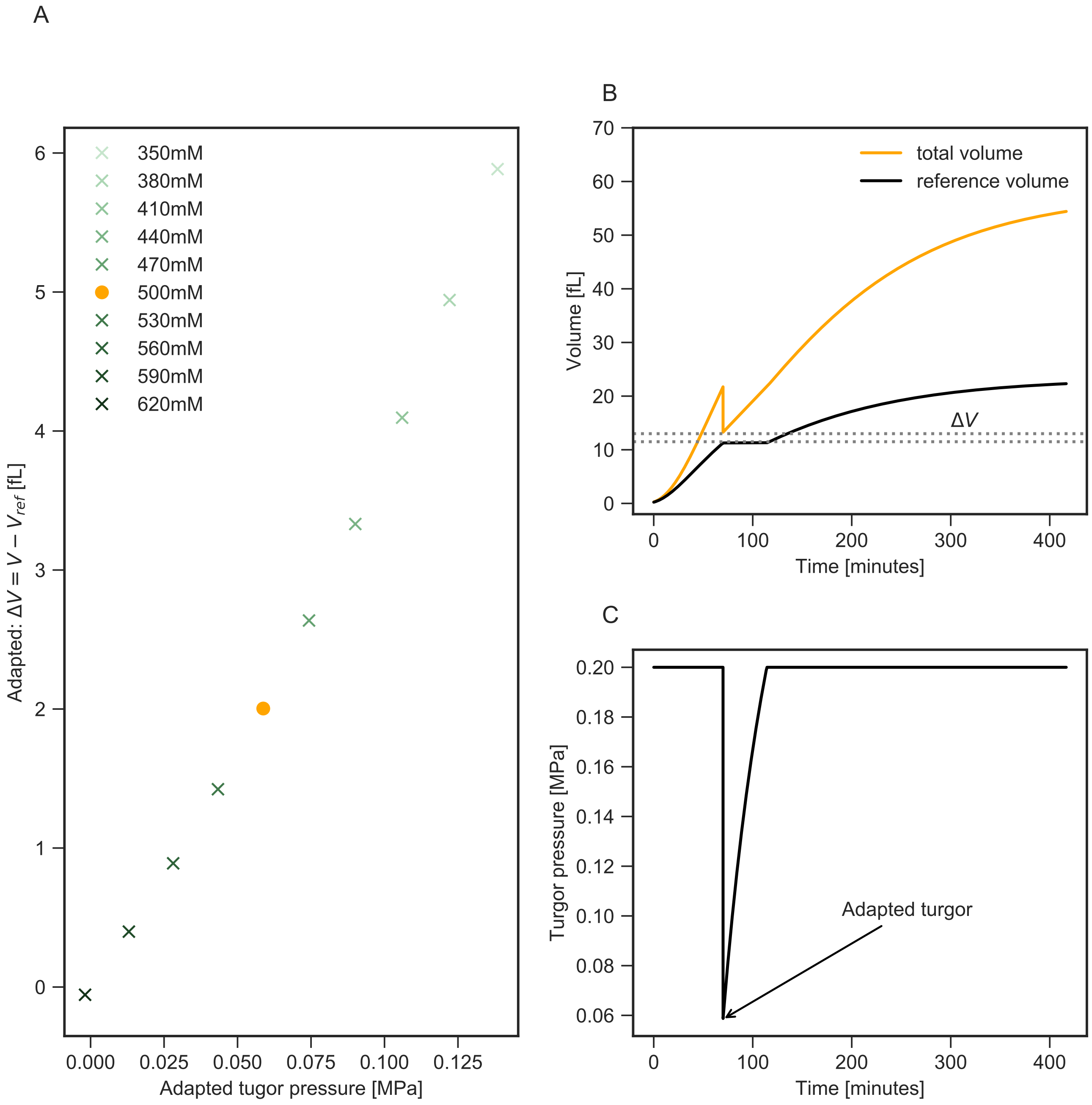


**Fig. 23. Total volume equals reference volume in case turgor pressure is zero, which proofs the proposed concept behind the reference volume of SCGM.** A: Each of the ten markers represents the outcome of a model simulation for ten different hyperosmotic shocks (light green to dark green: increasing external osmolarity). The highlighted dot (orange) represents the state, which follows a shock setting $\Pi_e = 500\ mM$ and is shown in detail in B and C. The total volume tends against the reference volume (immediate adaptation to the shock: difference between total volume and reference volume $V - V_{ref}$ becomes zero) in case turgor drops down to zero for most drastic shocks. B: illustrates the custom y-axis plotted in A and exemplify the simulation results of a shock setting the external osmolarity to $\Pi_e = 500\ mM$ (orange dot in A) by representatively showing the volume C: illustrates the custom x-axis of A and the turgor pressure response of the specific simulation indicated by the orange dot in A.

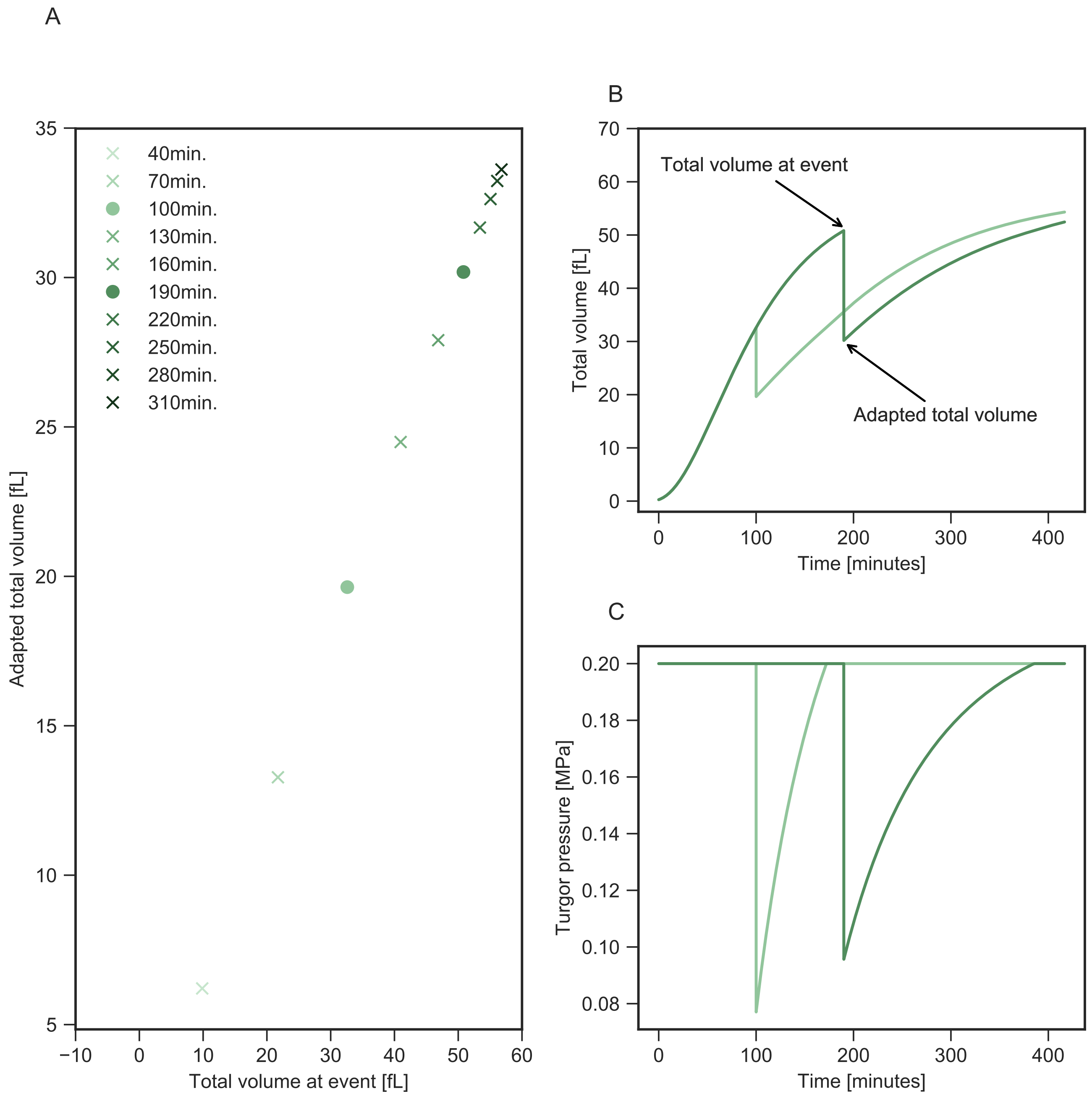


**Fig. 24. Total volume appearing due to a hyperosmotic shock is linearly related to absolute cell volume at event according to SCGM simulations.** A: Each of the ten markers represents the outcome of a model simulation for an equivalent shock but at ten different time points (light green to dark green: increasing time of shock event; shock was increasing external osmolarity from 240 mM to 500 mM in each case). The two filled circles (shocks at 100 minutes and 190 minutes) represent model simulations that are shown in detail in B and C. The total volume which adapts immediately after the shock (labeled adapted total volume and equal to the minimum volume following a shock) is linearly related to the size of the cell that experienced the shock (total volume at event). B: illustrates the custom x- and y-axis plotted in A and exemplifies two simulation results of equal shocks at the two time points 100 minutes and 190 minutes. Obviously, the two different time points link to two modeled cells of two distinct sizes but experience the same amount of shock. C: the turgor pressure response for an equal shock (from $c_e$=240 mM to 500 mM) for the two time points 100 minutes and 190 minutes (as indicated as filled circles in A).
